# PRMT5-mediated histone methylation regulates alternative splicing via MECP2-PTBP1 to promote EMT in breast cancer hypoxia

**DOI:** 10.1101/2024.10.16.618678

**Authors:** Srinivas Abhishek Mutnuru, Pooja Yadav, Parik Kakani, Shruti Ganesh Dhamdhere, Poorva Kumari, Shruti Agarwal, Atul Samaiya, Sanjeev Shukla

**Affiliations:** Department of Biological Sciences, Indian Institute of Science Education and Research Bhopal, Bhopal, Madhya Pradesh 462066, India; Beth Israel Deaconess Medical Center, Boston, MA 00215, U.S.A.; Department of Surgical Oncology, Bansal Hospital, Bhopal, Madhya Pradesh, 462016, India

**Keywords:** PRMT5, histone methylation, hypoxia, alternative splicing, EMT

## Abstract

Tumor hypoxia induced alterations in the epigenetic landscape and alternative splicing influence cellular adaptations. PRMT5 is a type II protein arginine methyltransferase that regulates several tumorigenic events in many cancer types. However, the regulation of PRMT5 and its direct implication on aberrant alternative splicing under hypoxia remains unexplored. In this study, we observed hypoxia induced upregulation of PRMT5 via the CCCTC binding factor, CTCF. Further, PRMT5-mediated symmetric arginine dimethylation H4R3me2s and H3R8me2s directly regulated the alternative splicing of Transcription Factor 3 (*TCF3*). Under hypoxia, PRMT5-mediated histone dimethylation at the intronic conserved region (ICR) present between *TCF3* exon 18a and exon 18b recruits DNA methyltransferase 3A (DNMT3A), resulting in DNA methylation. DNA methylation at the *TCF3*-ICR is recognized and bound by Methyl CpG binding protein (MECP2) resulting in RNA-Pol II pausing, promoting the recruitment of the negative splicing factor PTBP1 to the splicing locus of *TCF3* mRNA. PTBP1 promotes the exclusion of exon 18a which results in the production of the pro-invasive TCF3-18B (E47) isoform which promotes EMT and invasion of breast cancer cells under hypoxia. Collectively, our results indicate PRMT5-mediated symmetric arginine dimethylation of histones regulates alternative splicing of *TCF3* gene thereby enhancing EMT and invasion in breast cancer hypoxia.

## Introduction

Genesis of hypoxic niches in growing tumors triggers wide array of changes in tumor cells allowing them to circumvent the adverse effects of lack of oxygen (Andrysik et al., 2021; Z. Chen et al., 2023). Epigenetic reprogramming and regulation of alternative splicing events facilitates adaptability of cancer cells to challenging conditions(Z. Chen et al., 2023; Farina et al., 2020a; Kakani et al., 2024; Pandkar et al., 2023; Yadav et al., 2023). Epigenetic alterations and aberrant alternative splicing have been independently studied and shown to aid tumor cell progression under hypoxia. Considering the complexity and intricacies of alternative splicing, recent reports have suggested that chromatin structure, epigenetic marks on DNA and histone modifications regulate alternative splicing outcomes(Luco et al., 2010; Segelle et al., 2022a; Shukla et al., 2011). The role of epigenetic modifications in regulating alternative splicing is a new dimension which is currently drawing significant attention and requires deeper and better understanding. Specifically, the regulation of such interplay between epigenetic modifications and alternative splicing in the context of hypoxia remains significantly understudied.

Protein arginine methyl transferase 5 (PRMT5) catalyzes the symmetric dimethylation of arginine residues of both histone and non-histone proteins(Hwang et al., 2021; Kim & Ronai, 2020). PRMT5 has been reported to methylate histones H3 and H4 at R8 and R3 positions respectively. PRMT5 has been observed to be upregulated in various cancer types and is known to regulate crucial tumorigenic events such as proliferation, EMT, invasion and stem cell maintenance(Chiang et al., 2017; J. Gao et al., 2021; Jiang et al., 2018). Largely, PRMT5 has been reported to affect gene transcription via histone methylation and pre mRNA splicing via methylation of splicing factor proteins(Rengasamy et al., 2017; Sachamitr et al., 2021; Sengupta et al., 2021). Also, recent reports have suggested the role of hypoxia induced epigenetic and splicing aberrations in regulating tumor progression(Pandey et al., 2024; Yadav et al., 2023). However, there have been no studies so far which have explored the potential of PRMT5 in regulating alternative splicing via its histone methylation activity. Therefore, we aimed to bridge the void of knowledge and investigated the intriguing role of PRMT5-mediated histone methylation in regulating alternative splicing under hypoxia.

In this study, for the very first time, we have reported that PRMT5 is upregulated under hypoxia via CTCF, and its upregulation promotes EMT and invasion in hypoxic breast cancer cells. Globally, we observed that loss of PRMT5 majorly affected cassette exon events under hypoxia and we identified *TCF3* as one of the novel alternative splicing targets of PRMT5. Mechanistically we have shown that under hypoxia, PRMT5-mediated symmetric arginine dimethylation of histones H4 and H3 at R3 and R8 positions at an intronic conserved region (ICR) is directly responsible for regulation of alternative splicing of *TCF3*. Interestingly, we found that PRMT5-mediated histone methylation recruited DNMT3A to the splicing locus leading to DNA methylation of the *TCF3*-ICR region. Subsequently, we showed that binding of MeCP2 to the methylated DNA led to RNA pol II pausing under hypoxia which further resulted in the recruitment of the negative splicing factor PTBP1. PTBP1 helps in exon 18A exclusion under hypoxia resulting in the production of the TCF3-18B containing E47 isoform which helps tumor cell invasion under hypoxia. Notably, our study also addresses the intricate interplay between histone modifications, DNA methylation, RNA polymerase kinetics and splicing factor recruitment under hypoxia. This underlies the intimate dependency of chromatin modifications and splicing on each other and how they can be regulated in disease conditions to achieve adaptability. Remarkably, our findings for the first time, comprehensively demonstrate the nuanced relationship between PRMT5-mediated histone methylation, DNA methylation and alternative splicing governing tumor progression under hypoxia. This study opens new avenues for understanding hypoxia driven tumor progression through the interplay of PRMT5-mediated histone and DNA methylation, as well as alternative splicing. These regulatory networks could be potential targets for development of innovative therapeutic strategies against hypoxia inflicted breast tumors.

## RESULTS

### PRMT5 is upregulated in breast cancer cells under hypoxia

PRMT5 has been reported to be upregulated in several tumor types (Kim & Ronai, 2020; Sapir et al., 2021). However, the influence of hypoxia upon *PRMT5* gene regulation remains unexplored. To address this caveat, we first analyzed a publicly available patient microarray data obtained from TCGA (GSE76250) where in the samples were stratified as normoxic or hypoxic based on hypoxia signature gene expression. Among all the PRMTs, we found *PRMT5* to be the most significantly upregulated gene under hypoxia (Fig 1A-B). Additionally, we performed a Spearman’s correlation test to observe the correlation of *PRMT5* gene expression with 10 classical hypoxia gene markers (*P4HA1, ENO2, NEDD4L, HK2, LDHA, FOXO3, SLC2A1, BHLHE40, PGGK1, BNIP3L*) using TCGA breast cancer tumor gene sets (BRCA Tumor). We observed a significant positive correlation (R=0.42) between the gene expression of *PRMT5* and hypoxia markers (Fig 1C) indicating that hypoxia regulates *PRMT5* gene expression in human breast cancers. Next, to validate *PRMT5* expression under hypoxia, we performed qRT-PCR and immunoblotting to understand *PRMT5* gene and protein expression in two breast cancer cell lines MCF7 and MDA-MB-231 subjected to hypoxia. We found a significant upregulation in *PRMT5* gene (Fig 1D) and protein expression (Fig 1E) under hypoxia in both MCF7 and MDA-MB231 breast cancer cell lines. PRMT5 symmetrically dimethylates H3R8 and H4R3. Hence, we performed immunoblotting analysis to check the expression of PRMT5-mediated histone modifications H3R8me2s and H4R3me2s under hypoxia. We observed an increase in PRMT5-mediated histone marks H3R8me2s and H4R3me2s under hypoxia in both MCF7 and MDA-MB-231 cell lines (Fig 1E) indicating that PRMT5 upregulation could have functional implication via histone modifications under hypoxia. To extend our observations to clinical samples, we checked the *PRMT5* transcript and protein levels along with the PRMT5-mediated modifications H4R3me2s and H3R8me2s in patient derived breast cancer cell line BC8322, subjected to hypoxia. We observed a significant increase in *PRMT5* mRNA (Fig 1F) as well as protein expression along with the corresponding increase in histone marks H4R3me2s and H3R8me2s under hypoxia (Fig 1G). Next, we performed IHC-F to detect PRMT5 expression in tumor sections derived from breast cancer patients. CA9 was used as a marker for hypoxia positive tumor regions. Our immunostaining analysis showed a positive correspondence of CA9 and PRMT5 expression. Evidently, we found that regions with negative CA9 staining, indicative of non-hypoxic regions, showed reduced PRMT5 expression (Fig 1H, S1A). We also observed a significant positive correlation between PRMT5 and CA9 fluorescent staining indicating that PRMT5 is upregulated under hypoxia (Fig. 1I) suggesting that PRMT5 upregulation is clinically relevant and can have implications in governing tumor progression. Together, these results indicate that PRMT5 is upregulated in breast cancer cells under hypoxia.

**Fig 1.**
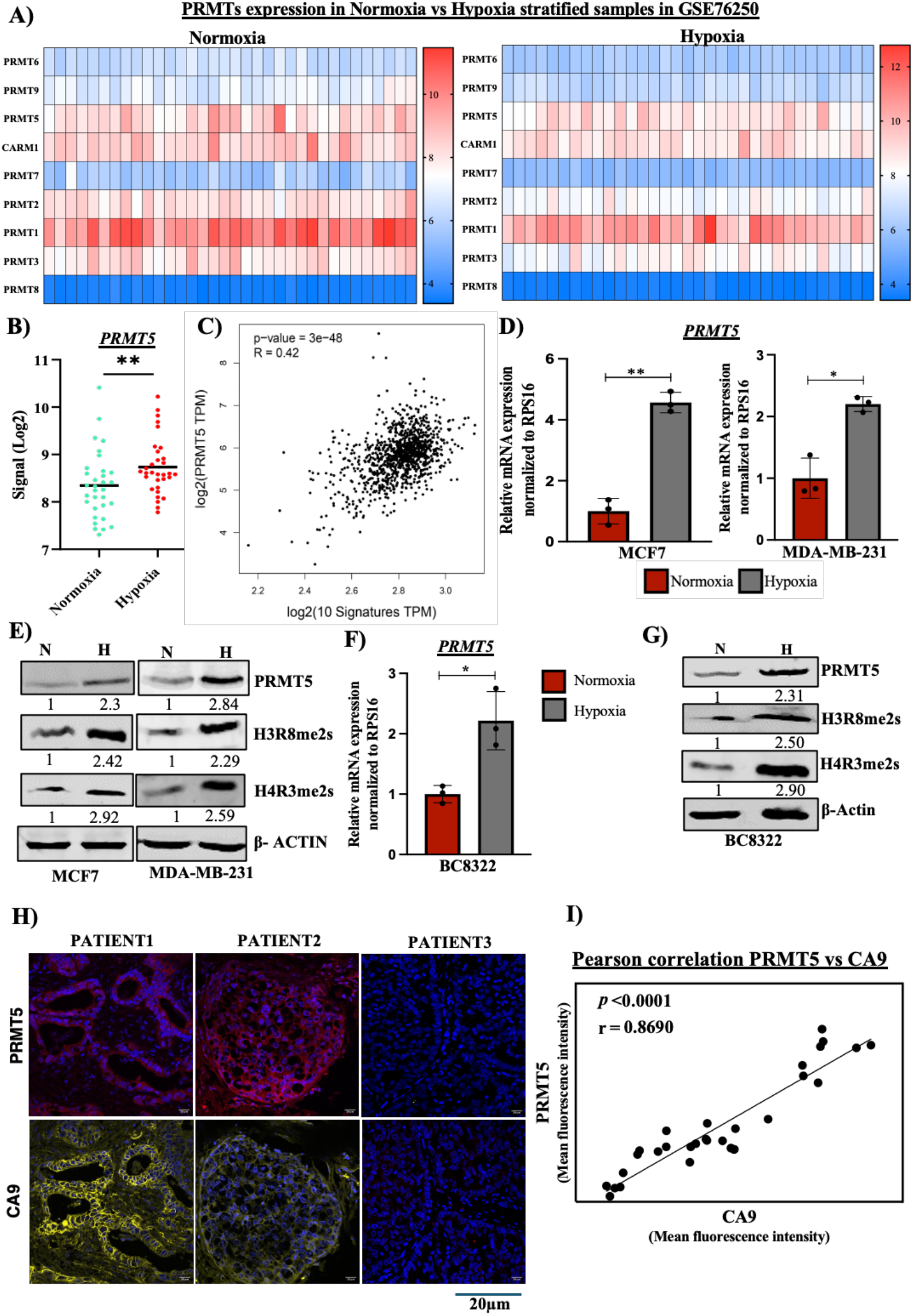
A) Heat maps showing Log2 signals of all PRMTs Normoxia vs Hypoxia in breast cancer patient microarray (GSE76520) stratified as normoxia or hypoxia. B) PRMT5 mRNA expression in breast cancer patient microarray (GSE76520) stratified as normoxia or hypoxia. C) Spearman’s correlation analysis of PRMT5 mRNA expression with 10 hypoxia marker genes. D) *PRMT5* mRNA expression in MCF7 and MDA-MB-231 breast cancer cell lines subjected to hypoxia. E) Immunoblot showing PRMT5, H4R3me2s and H3R8me2s expression under hypoxia in MCF7 and MDA-MB-231 cell lines. F) *PRMT5* mRNA expression in patient derived cell line BC8322 subjected to hypoxia. G) Immunoblot showing PRMT5, H4R3me2s and H3R8me2s expression under hypoxia in in patient derived cell line BC8322. H) IHC-F analysis showing PRMT5 and CA9 expression in tumor sections of breast cancer patients. I) Pearson correlation analysis between PRMT5 and CA9 fluorescence intensities calculated from IHC-F. Scale bar 20μm. Error bars, mean ± SEM; two-tailed t test, ∗*p* < 0.05, ∗∗*p* < 0.01, ∗∗∗*p* < 0.001, n = 3 biological replicates.

### CTCF is responsible for PRMT5 upregulation under hypoxia

Following our observation that PRMT5 is upregulated under hypoxia, we next wanted to identify the factors responsible for transcriptional regulation of *PRMT5* under hypoxia. Since we observed a significant increase in *PRMT5* mRNA expression, we hypothesized that *PRMT5* is transcriptionally regulated under hypoxia. We generated 4 sequential promoter deletion constructs spanning the *PRMT5* promoter up to -2000bp upstream of the TSS with 500bp deletions in each construct. These promoter constructs were cloned upstream of firefly luciferase (Fluc) gene in a pGL3-basic vector (Fig 2A). The luciferase construct harboring the full-length promoter (+1 to -2000bp) segment and the 1500 bp segment (+1 to -1500bp) upstream of transcription start site showed a significant increase in luciferase activity under hypoxia in both MCF7 and MDA-MB-231 cell lines (Fig. 2B, S2A). This observation was indicative of the fact that a regulatory region present between -1000 to -2000 bp of the *PRMT5* promoter was responsible for regulating *PRMT5* expression under hypoxia. We then made use of publicly available chip-seq datasets from ChIP-Atlas to understand which transcription factors might potentially bind and regulate *PRMT5* expression under hypoxia. We identified a significant CTCF binding site within the regulatory region at -1175bp position of the promoter which also had a significant binding score in the analysis performed in JASPAR (Fig S2B). Following this, we verified CTCF binding at *PRMT5* promoter under hypoxia using ChIP-Seq data in MCF7 cell line (Kakani et al., 2023)(Fig 2C).

**Fig 2.**
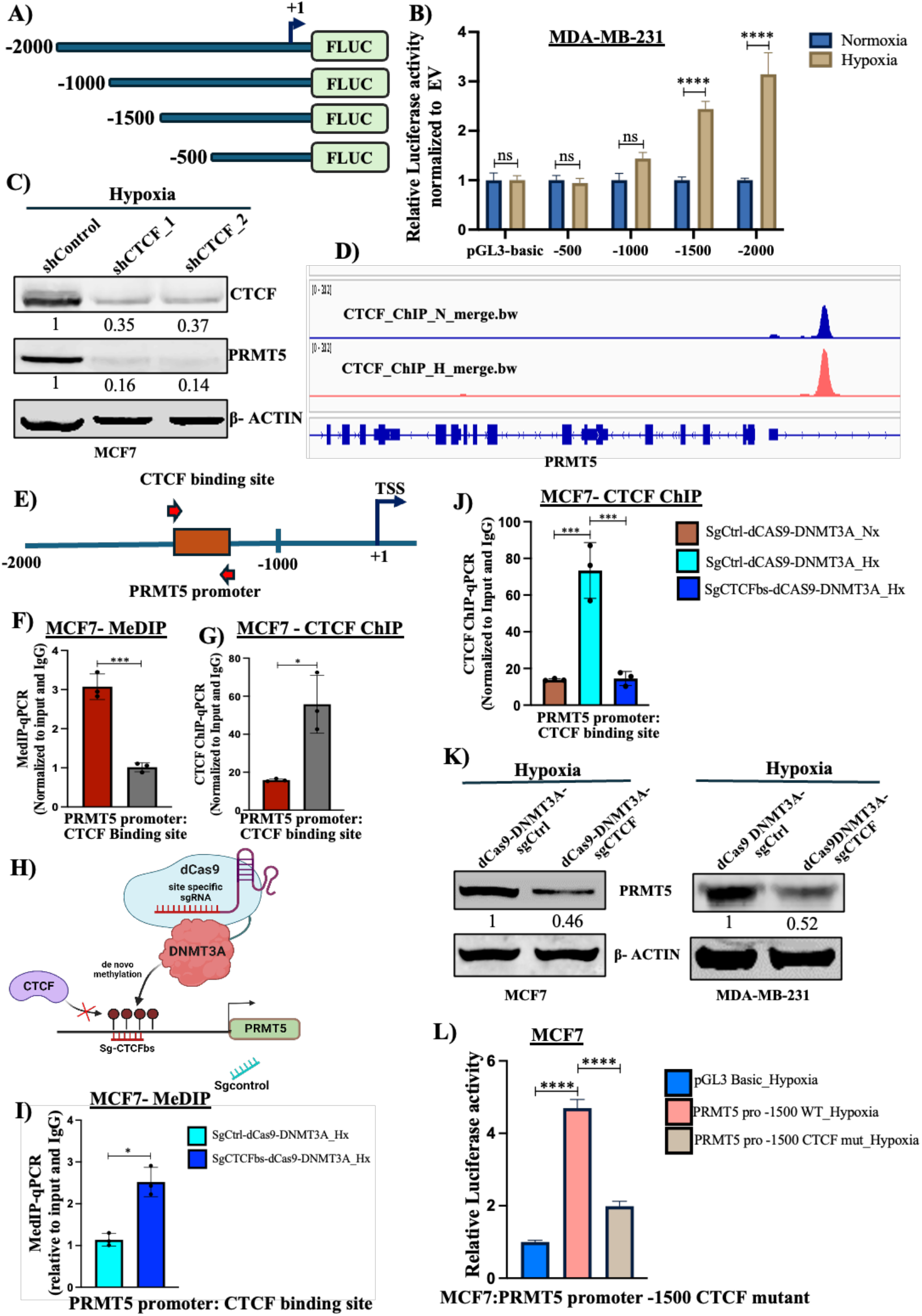
A) Representation of PRMT5 promoter luciferase construct cloned upstream of F-Luc gene in pGL3 basic vector. B) Luciferase assay showing an increase in luciferase activity under hypoxia in MDA-MB-231 cells. C) Immunoblot showing decrease in PRMT5 expression upon CTCF KD under hypoxia in MCF7 cells. D) MCF7 CTCF ChIP-seq track showing CTCF enrichment at PRMT5 promoter under hypoxia. E) Schematic representation of CTCF binding site at PRMT5 promoters along with the primer positions. F) MeDIP-qPCR showing decrease in DNA methylation at PRMT5 promoter in MCF7 cells normoxia vs hypoxia. G) CTCF Chip qPCR showing enrichment in CTCF binding at PRMT5 promoter in MCF7 cells normoxia vs hypoxia. H) Schematic representation of the dCAS9-DNMT3A epigenetic system targeting CTCF binding site of PRMT5 promoter. I) MeDIP qPCR showing enrichment of DNA methylation at PRMT5 promoter under hypoxia post dCAS9-DNMT3A sgCTCFbs vs sgControl transfections. J) CTCF ChIP qPCR showing reduction in CTCF binding at PRMT5 promoter under hypoxia post dCAS9-DNMT3A sgCTCFbs vs sgControl transfection. K) Immunoblot showing decrease in PRMT5 expression upon transfection with dCAS9-DNMT3A sgCTCFbs vs sgControl. L) Luciferase assay showing decrease in luciferase activity in promoter construct harboring a mutated CTCF binding site. Error bars, mean ± SEM; two-tailed t test, one way ANOVA. ∗*p* < 0.05, ∗∗*p* < 0.01, ∗∗∗*p* < 0.001, n = 3 biological replicates.

Firstly, to establish the role of CTCF in regulating *PRMT5* gene expression under hypoxia, we performed shRNA mediated CTCF knock down in both MCF7 and MDA-MB-231 cells. Our immunoblotting analysis revealed a significant reduction in PRMT5 protein expression upon CTCF knock down under hypoxia in both MCF7 and MDA-MB-231 cell lines (Fig 2D, S2C). DNA methylation is known to regulate gene expression and since CTCF binding is also sensitive to DNA methylation, we performed MeDIP-qRT-PCR to analyze the DNA methylation profile at the CTCF binding site of the *PRMT5* promoter (Fig 2E) to understand the role of promoter DNA methylation in regulating *PRMT5* expression under hypoxia. We observed a significant decrease in DNA methylation at the CTCF binding site of the *PRMT5* promoter (Fig 2F, S2D) under hypoxia indicating that reduced DNA methylation under hypoxia could possibly allow the binding of CTCF to regulate *PRMT5* gene expression. Next, to confirm the binding of CTCF at the *PRMT5* promoter, we performed CTCF ChIP-qPCR assay and observed a significant increase in CTCF occupancy at the CTCF binding site of the *PRMT5* promoter in both MCF7 and MDA-MB-231 cell lines (Fig 2G, S2E) suggesting that reduced DNA methylation at the CTCF binding site allows CTCF binding on *PRMT5* promoter under hypoxia. In order to further verify the role of DNA methylation in regulating *PRMT5* expression under hypoxia, we made use of the dCAS9-DNMT3A epigenetic modifier system to preferentially methylate the CTCF binding site of the *PRMT5* promoter (Fig 2H). The dCAS9-DNMT3A construct with the CTCF binding site targeting sgRNA was transfected into MCF7 and MDA-MB-231 cells under hypoxia following which we analyzed for DNA methylation profile, CTCF binding and *PRMT5* expression. Firstly, we observed a significant increase in DNA methylation (Fig 2I) and a concomitant decrease in CTCF binding at the CTCF binding site (Fig. 2J) of the *PRMT5* promoter upon transfection with the dCAS9-DNMT3A construct targeting the CTCF binding site of the *PRMT5* promoter under hypoxia, suggesting that decrease in methylation under hypoxia is necessary for CTCF binding at the *PRMT5* promoter. Expectantly, our immunoblot analysis revealed a significant decrease in PRMT5 protein expression (Fig. 2K) indicating that under hypoxia, decrease in DNA methylation allows CTCF binding at the *PRMT5* promoter thereby regulating *PRMT5* expression. Furthermore, we mutated the CTCF binding site present in the promoter-luciferase construct (+1 to -1500bp) (Fig S2F) and performed luciferase assay under hypoxia to establish the role of CTCF mediated transcriptional upregulation of *PRMT5*. In corroboration with our previous results, we observed a significant reduction in luciferase activity in the construct harboring mutant CTCF binding site under hypoxia (Fig 2L, S2G) suggesting that CTCF is responsible for *PRMT5* upregulation under hypoxia. Put together, our observations suggest that reduction in DNA methylation at the CTCF binding site of the *PRMT5* promoter allows CTCF binding and regulation of *PRMT5* expression under hypoxia.

### PRMT5 regulates EMT and invasion of breast cancer cells under hypoxia

Metastasis is responsible for more than 90% of cancer related mortalities(Chaffer & Weinberg, 2011; Gupta & Massagué, 2006; Riggio et al., 2021). Therefore, it becomes imperative to improve our understanding of the molecular mechanisms governing the complex process of metastatic cascade in order to tackle its adverse effects. Epithelial to Mesenchymal Transition (EMT) is a process wherein the epithelial cells acquire characteristic features of mesenchymal cells conferring epithelial cells the ability to invade and metastasize. PRMT5 is known to regulate EMT and invasion in different cancer types(H. Chen et al., 2017; L. Ge et al., 2020; J. Huang et al., 2021). Also, hypoxia is known to induce EMT and invasion of cancer cells(Kakani et al., 2024). However, there have been no reports with respect to the role of PRMT5 in regulation of EMT and invasion under hypoxia. Therefore, we envisaged to understand the role of PRMT5 in regulating EMT and invasion under hypoxia. Firstly, we performed shRNA mediated PRMT5 knockdown under hypoxia in both MCF7 and MDA-MB-231 cell lines to check the effect of PRMT5 on EMT induction. We performed q-RT-PCR to check the transcript levels of the mesenchymal marker genes *VIM* (Vimentin) and *CDH2* (N-Cadherin) along with the epithelial marker gene *CDH1* (E-Cadherin) and the EMT regulating transcription factor *SNA1* (Snail) in both MCF7 and MDA-MB-231 cell lines. We observed a significant decrease in the transcript levels of the mesenchymal markers *VIM* and *CDH2* along with EMT transcription factor *SNAI1* (Fig. 3A, S3A). Epithelial marker CDH1 showed a significant increase in transcript levels upon PRMT5 KD under hypoxia in both MCF7 and MDA-MB-231 cell lines (Fig. 3A, S3A). We then analyzed the protein levels of epithelial markers E-cad and Cytokeratin-18 in MCF7, and Cytokeratin-18 in MDA-MB-231. Protein levels of mesenchymal marker Vimentin along with EMT regulating transcription factor Snail were also analyzed in both MCF7 and MDA-MB-231 cell lines. Our immunoblot analyses showed an increase in the epithelial marker E-Cad and Cytokeratin 18 upon PRMT5 knockdown under hypoxia in both MCF7 and MDA-MB-231 cell lines whereas the mesenchymal marker Vimentin along with the transcription factor Snail showed decreased protein expression upon PRMT5 knockdown under hypoxia in both MCF7 and MDA-MB-231 cell lines (Fig. 3B, S3B). Therefore, these results suggested that PRMT5 somehow regulated EMT induction under hypoxia.

**Fig 3.**
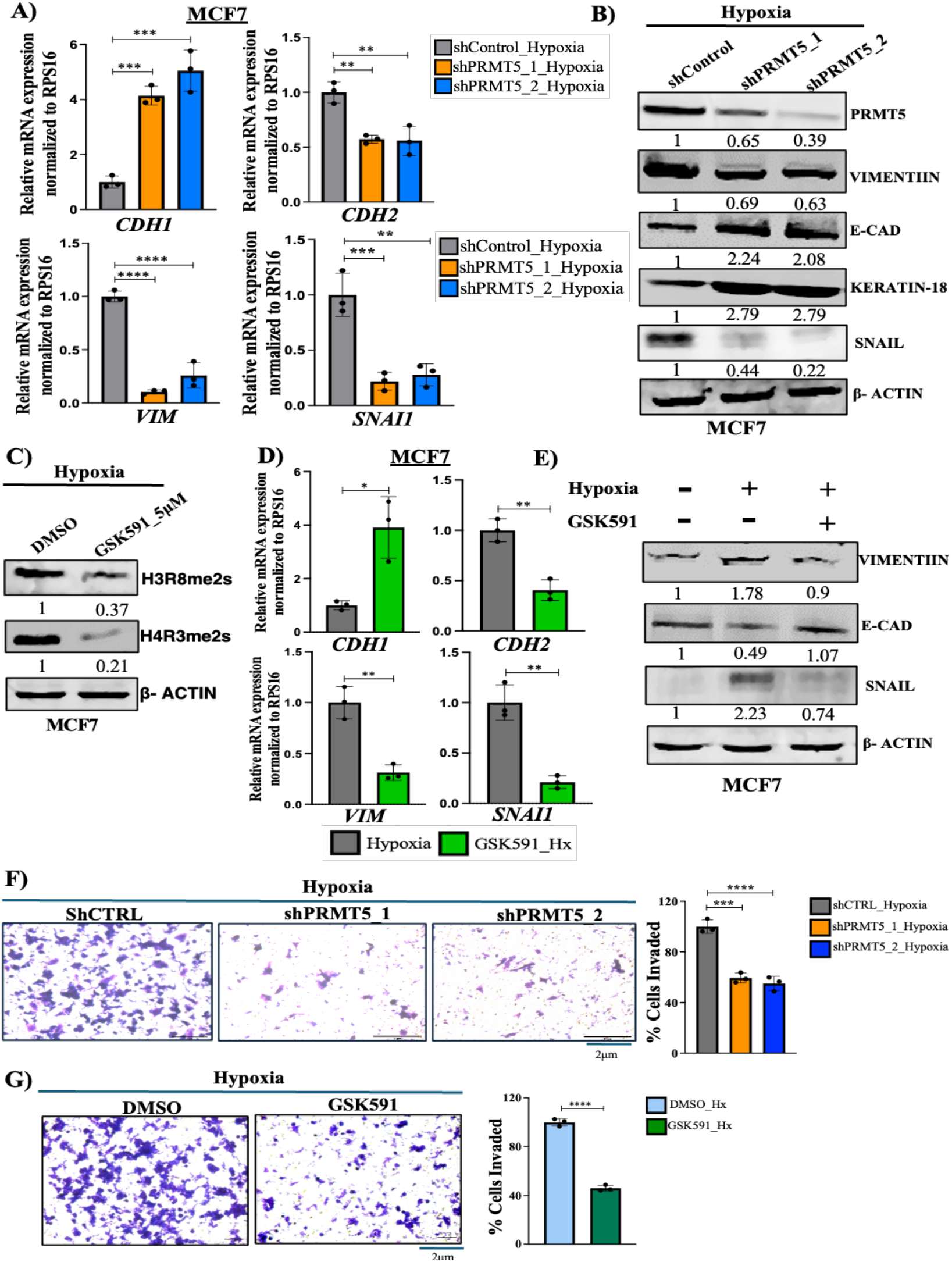
A) qRT-PCR depicting mRNA expression of EMT markers in shCTRL vs shPRMT5 MCF7 cells under hypoxia. B) Immunoblot showing protein level of EMT markers in shCTRL vs shPRMT5 MCF7 cells under hypoxia. C) Immunoblot showing decrease in histone marks HR38me2s and H4R3me2s upon vehicle (DMSO) vs 5μM GSK591 treatment in MCF7 cells under hypoxia. D) qRT-PCR depicting mRNA expression of EMT markers upon vehicle (DMSO) vs 5μM GSK591 treatment in MCF7 cells under hypoxia. E) Immunoblot showing protein level of EMT markers upon vehicle (DMSO) vs 5μM GSK591 treatment in MCF7 cells under hypoxia. F) Matrigel invasion assay and its respective quantification (right side) in shCTRL vs shPRMT5 PRMT5 KD MCF7 cells under hypoxia G) Matrigel invasion assay and its respective quantification (right side) upon vehicle (DMSO) vs 5μM GSK591 treatment in MCF7 cells under hypoxia. Scale bar 2μm. Error bars, mean ± SEM; two-tailed t test, one way ANOVA. ∗*p* < 0.05, ∗∗*p* < 0.01, ∗∗∗*p* < 0.001, n = 3 biological replicates.

As observed earlier, PRMT5-mediated histone modifications H3R8me2s and H4R3me2s were found to be upregulated under hypoxia. Therefore, we hypothesized if obliterating these histone modifications was sufficient to curb EMT induction under hypoxia. To test this, we used GSK591 which is a substrate competitive inhibitor of PRMT5 that exerts its effect by blocking the ability of PRMT5-MEP50 complex to methylate target proteins(Chan-Penebre et al., 2015; Duncan et al., 2016). We treated MCF7 and MDA-MB-231 cells with 5μM GSK591 under hypoxia and found a significant reduction in the histone marks H3R8me2s and H4R3me2s (Fig. 3C, S3C) indicating that GSK591 specifically inhibits PRMT5 activity. Along with this, as observed earlier, our q-RT-PCR results showed a significant decrease in the transcript levels of the mesenchymal markers *VIM* and *CDH2* along with EMT transcription factor *SNAI1* whereas the Epithelial marker *CDH1* showed a significant increase in transcript levels (Fig. 3D, S3D). We also found that the epithelial markers E-cad (MCF7) and Cytokeratin 18 (MDA-MB-231) showed an increase in protein expression upon PRMT5 activity inhibition under hypoxia, whereas the mesenchymal marker Vimentin along with the transcription factor Snail showed decreased protein expression upon PRMT5 activity inhibition under hypoxia in MCF7 and MDA-MB-231 cell lines (Fig. 3E, S3E). Together, these observations suggested that PRMT5-mediated histone modifications (H3R8me2s/H4R3me2s) were involved in regulation of EMT induction under hypoxia.

Induction of EMT subsequently renders epithelial cells more invasive which initially invade through the adjacent basement membrane to initiate the first step of the metastatic cascade (Cao et al., 2023). Therefore, we intended to study PRMT5-mediated acquisition of invasive potential of breast cancer cells under hypoxia. We performed Matrigel invasion assay and observed a significant decrease in the invasive potential of both MCF7 and MDA-MB-231 cell lines upon PRMT5 knock down under hypoxia (Fig. 3F, S3F). Additionally, treatment of MCF7 and MDA-MB-231 cell lines with 5μM GSK591 similarly decreased the invasive potential of both breast cancer cell lines under hypoxia (Fig. 3G, S3G). In summary, our findings demonstrated that PRMT5-mediated histone modifications regulate EMT and invasion of breast cancer cells under hypoxia.

### PRMT5 regulates *TCF3* splicing under hypoxia

So far, we established that PRMT5-mediated histone modifications regulated EMT and invasion under hypoxia. Next, we wanted to delineate the mechanism by which PRMT5 regulated EMT under hypoxia via modulating the histone marks. PRMT5-mediated symmetric arginine dimethylation of histones influences chromatin structure and accessibility thereby governing gene transcription(Q. Gao et al., 2023; Liu et al., 2020). Additionally, PRMT5 is also known to regulate the process of alternative splicing by methylating the essential components of the spliceosome complex (Rengasamy et al., 2017). Hypoxia also triggers aberrant alternative splicing patterns aiding tumor progression (Farina et al., 2020b; Yadav et al., 2023). Lately, numerous studies have shown that owing to the co-transcriptional nature of splicing, changes in the epigenetic landscape including histone and DNA modifications intricately regulate alternative splicing outcomes(Pandey et al., 2024; Segelle et al., 2022b; Shukla et al., 2011; Yadav et al., 2023). Notably, despite the aforementioned insights, there have been no reports thus far which show how PRMT5 regulates alternative splicing under hypoxia via modulation of histone methylation. Therefore, we set out to understand PRMT5-mediated regulation of alternative splicing via modulation of histone modifications under hypoxia.

Firstly, to understand the effect of PRMT5-mediated changes in alternative splicing events under hypoxia, we performed shRNA mediated knock down of PRMT5 under hypoxia in MDA-MB-231 cells followed by RNA-Seq. RNA-Seq analysis was performed and various splicing events were classified. Among the total significant splicing events identified (p<0.05), 67% were cassette exon events (skipped exons), 15% were mutually exclusive exons (MXE), 4% were retained introns (RI), and 8% and 6% were alternative 5’ splice site (A5SS) and alternative 3’ splice site (A3SS) selection events, respectively (Fig. 4A). Since PRMT5 KD majorly led to changes in cassette exon events, we further analyzed the cassette exon events and selected only those events which had |®PSI| > 3% and p<0.05. Among the significant cassette exon events, we found 59.35% events to be exon inclusion events and 40.46% to be exon exclusion events (Fig. 4B). Since exon inclusion was the major alternative splicing event type upon PRMT5 KD under hypoxia, we further analyzed exon inclusion events. Gene Ontology (GO) analysis of the genes showing exon inclusion upon PRMT5 knockdown under hypoxia revealed regulation of apical junctions and epithelial-mesenchymal transition (EMT) to be among the top enriched categories (Fig. 4C). This further suggested that PRMT5-mediated changes in alternative splicing significantly impacted EMT processes under hypoxia.

**Fig 4.**
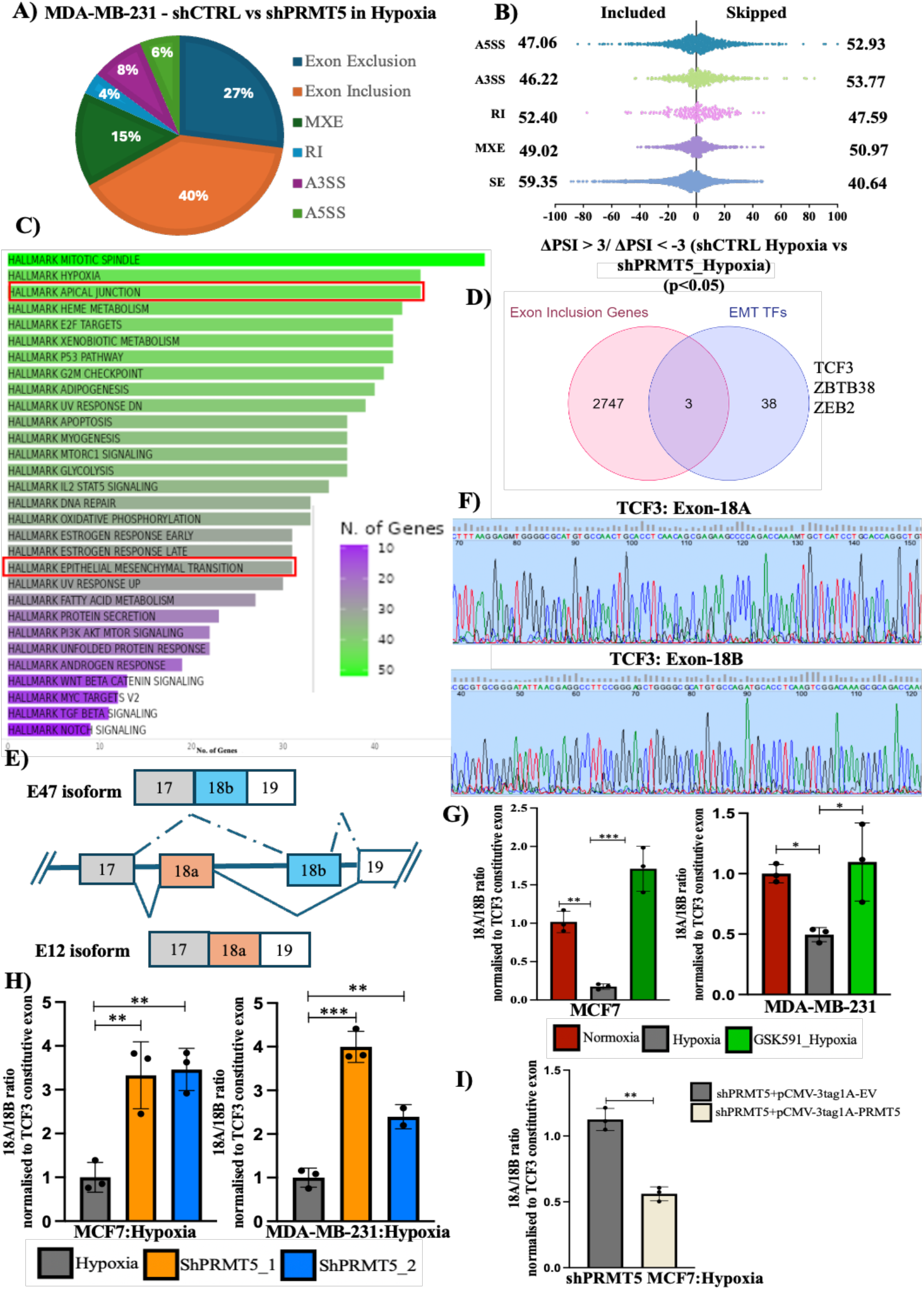
A) Pie chart showing distribution of different types of significant AS events (p<0.05) in shCTRL vs shPRMT5 MDA-MB-231 cells under hypoxia. B) Distribution of ®PSI of significant AS events in shCTRL vs shPRMT5 MDA-MB-231 cells under hypoxia. C) Gene ontology analysis of genes showing significant exon inclusion upon PRMT5 KD under hypoxia. D) Venn diagram showing *TCF3, ZBTB38* and *ZEB2*, as novel alternatively spliced targets of PRMT5 obtained by overlapping EMT transcription factors gene set (ref) vs gens having significant exon inclusion changes in shCTRL vs shPRMT5 MDA-MB-231 cells under hypoxia. E) Schematic depiction of *TCF3* alternative splicing. F) Chromatogram showing PCR products obtained using TCF3-18A and TCF3-18B exon specific primers. G) qRT-PCR showing change in exon 18A/18B ratio upon vehicle (DMSO) vs 5μM GSK591 treatment in MCF7 and MDA-MB-231 cells H) qRT-PCR showing an increase in exon 18A/18B ratio in shCTRL vs shPRMT5 MCF7 and MDA-MB-231 cells under hypoxia. I) qRT-PCR showing a decrease in 18A/18B ratio upon PRMT5 OE in shPRMT5 MCF7 cells under hypoxia. Error bars, mean ± SEM; two-tailed t test, one way ANOVA. ∗*p* < 0.05, ∗∗*p* < 0.01, ∗∗∗*p* < 0.001, n = 3 biological replicates.

EMT transcription factors are critical regulators of EMT induction and are essential for the initiation and progression of cancer cells to a more aggressive and metastatic phenotype(Y. Huang et al., 2022; Imani et al., 2016; Saitoh, 2023). Therefore, we overlapped our dataset of genes showing exon inclusion events with a dataset of EMT transcription factors (Debnath et al., 2021) with the aim to identify EMT factors undergoing PRMT5-mediated alternative splicing under hypoxia. Among all the EMT regulating transcription factors, TCF3, ZEB2 and ZBTB38 showed significant exon inclusion changes upon PRMT5 KD under hypoxia (Fig.4D). To delineate the detailed mechanism of how PRMT5 regulated alternative splicing under hypoxia we further selected *TCF3* as our model gene system.

Transcription Factor 3 (TCF3) is a member of the basic-helix-loop-helix (bHLH) transcription factors that has been previously shown to repress E-cad expression in differentiated cells thereby promoting invasive phenotype(Yamazaki et al., 2018). Additionally, *TCF3* gene has been shown to promote cancer invasiveness and stemness in different cancer types(L. Li et al., 2019; Z. X. Li et al., 2023; López-Menéndez et al., 2021). So far, there have been no reports suggesting how *TCF3* splicing is affected in tumor hypoxia and its role in orchestrating EMT and invasion. Therefore, we hypothesized that PRMT5 regulated *TCF3* splicing via modulating the histone methylation thereby promoting EMT and invasion under hypoxia.

*TCF3* alternative splicing event involves exon 18A and exon 18B. Inclusion of exon 18A results in the production of the TCF3-18A (E12) isoform whereas, inclusion of 18B leads to the production of the pro-invasive TCF3-18B (E47) isoform(Yamazaki et al., 2018). Our RNA-Seq analysis displayed increased inclusion of exon 18A (|®PSI| > 3% and p<0.05) upon PRMT5 KD under hypoxia. Since these exons are mutually included or excluded in the final transcript, we wanted to analyze how exon 18A inclusion was regulated with respect to exon 18B under hypoxia. Firstly, to understand *TCF3* splicing pattern and its dependency on PRMT5 in breast cancer hypoxia, we performed qRT-PCR analysis with or without GSK591 treatment using primers designed specifically against exons 18A and 18B. We sequence verified the specificity of these primers and confirmed that they amplified the intended exons only (Fig. 4F). We observed that, under hypoxia, *TCF3* exon 18A/18B ratio was significantly downregulated but was rescued back upon PRMT5 activity inhibition by GSK591 indicating that exon 18A inclusion was significantly downregulated under hypoxia in both MCF7 and MDA-MB-231 cells and PRMT5 was involved in exon 18A exclusion under hypoxia (Fig. 4G). Next, we performed shRNA mediated knockdown of PRMT5 in both MCF7 and MDA-MB-231 cell lines under hypoxia to verify PRMT5-mediated regulation of *TCF3* splicing. Our qRT-PCR analysis showed an increase in exon 18A/18B ration upon PRMT5 knockdown under hypoxia (Fig. 4H). To rescue this effect, we performed shRNA mediated knock down of PRMT5 under hypoxia in MCF7 cells in which we over expressed PRMT5 (Fig. 4I) following which we performed qRT-PCR analysis to check if replenishing PRMT5 expression overturned the splicing event. As expected, over expression of PRMT5 under hypoxia led to a significant decrease in the exon 18A/18B ratio indicating that PRMT5 somehow restricted 18A inclusion under hypoxia (Fig.4J). Together, these results showed that PRMT5-mediated symmetric arginine histone dimethylation regulates *TCF3* splicing event under hypoxia by inhibition of exon 18A inclusion.

### PRMT5-mediated H4R3me2s modification is essential and sufficient to regulate *TCF3* splicing under hypoxia

Until now, we delineated that PRMT5-mediated histone methylation regulated *TCF3* splicing under hypoxia. We next wanted to systematically deduce the mechanism of how the histone methylation performed by PRMT5 affected the *TCF3* splicing outcome under hypoxia. The *TCF3* gene harbors an intronic conserved region (ICR) which has already been reported to be a site of alternative splicing regulation concerning exons 18A and 18B(Yamazaki et al., 2019). Therefore, we investigated if this could potentially be a site which could be utilized by PRMT5 for exerting its effect on *TCF3* splicing regulation. To check this, we performed H3R8me2s and H4R3me2s ChIP-qPCR analysis at the *TCF3*-ICR region in both MCF7 and MDA-MB-231 cells under hypoxia and observed a significant increase in both the histone marks H3R8me2s and H4R3me2s at the *TCF3*-ICR region (Fig. 5B, S4A).

**Fig 5.**
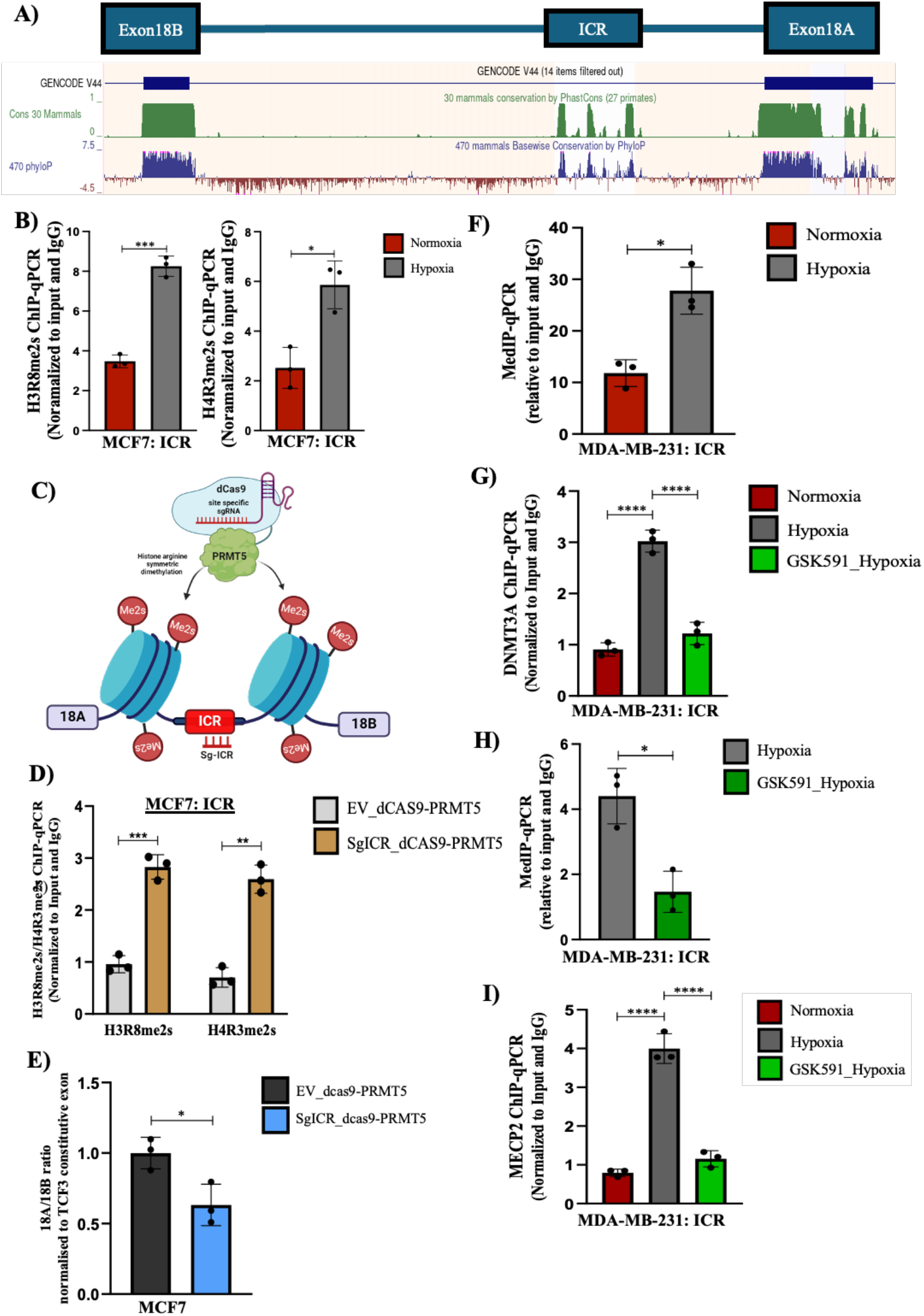
A) Genome track from UCSC genome browser showing TCF3 Intronic Conserved Region (ICR) B) H3R8me2s and H4R3me2s ChIP qPCR depicting increased histone symmetric arginine dimethylation at TCF3-ICR region in MCF7 cells, normoxia vs hypoxia. C) Schematic representation of the dCAS9-PRMT epigenome editing vector targeted at the TCF3-ICR region D) H3R8me2s and H4R3me2s ChIP qPCR depicting increased histone symmetric arginine dimethylation at TCF3-ICR region after transfection with dCAS9-PRMT5-EV vs dCAS9-PRMT5-sgICR in MCF7 shPRMT5 cells. E) qRT-PCR showing decrease in exon 18A/18B ratio after transfection with dCAS9-PRMT5-EV vs dCAS9-PRMT5-sgICR in MCF7 shPRMT5 cells. F) MeDIP qPCR showing increase in DNA methylation at TCF3-ICR region in MDA-MB-231 cells, normoxia vs hypoxia. G) DNMT3A ChIP qPCR showing change in DNMT3A binding at TCF3-ICR region in MDA-MB-231 cells treated with DMSO (Nx vs Hx) or 5μM GSK591(Hx). H) MeDIP qPCR showing decrease in DNA methylation at TCF3-ICR region in MDA-MB-231 cells after treatment with DMSO vs 5μM GSK591under hypoxia. I) MECP2 ChIP qPCR depicting change in MECP2 binding at TCF3-ICR region in MDA-MB-231 cells treated with DMSO (Nx vs Hx) or 5μM GSK591(Hx). Error bars, mean ± SEM; two-tailed t test, one way ANOVA. ∗*p* < 0.05, ∗∗*p* < 0.01, ∗∗∗*p* < 0.001, n = 3 biological replicates.

To establish that PRMT5-mediated histone methylation at the *TCF3*-ICR region is essential to regulate alternative splicing we made use of a dCAS9 epigenome modification system. We, for the first time generated a dCAS9-PRMT5 construct containing the full length PRMT5 gene downstream of dCAS9 which can be used to regulate symmetric dimethylation of histones at target regions (Fig S4B-C). We made use of this dCAS9-PRMT5 construct to target the *TCF3*-ICR region and checked the effect of local histone symmetric dimethylation event on *TCF3* splicing (Fig. 5C). Firstly, we performed shRNA mediated knockdown of PRMT5 to deplete endogenous PRMT5 protein following which the cells were transfected with the dCAS9-PRMT5 construct targeting the *TCF3*-ICR region to specifically modify the local histone symmetric dimethylation mark. We then verified if the dCAS9-PRMT5 construct was able to induce symmetric dimethylation of histones at H4R3 and H3R8 positions at the desired position at the splicing locus. Our ChIP-qPCR analysis showed that targeting dCAS9-PRMT5 at the *TCF3*-ICR region was able to catalyze both H4R3me2s and H3R8me2s modifications at the *TCF3*-ICR region (Fig. 5D). We then used the construct to check if local modifications of H4R3me2s and H3R8me2s at the *TCF3*-ICR region was sufficient to alter *TCF3* splicing event. We performed qRT-PCR analysis following dCAS9-PRMT5 transfection in PRMT5 knockdown MCF7 cells to check the change in *TCF3* alternative splicing pattern. We observed a significant decrease in the 18A/18B ratio upon targeting dCAS9-PRMT5 to the *TCF3* splicing locus (Fig. 5E) indicating that local histone arginine symmetric dimethylation catalyzed by PRMT5 at the *TCF3* splicing locus is sufficient to alter the alternative splicing pattern of *TCF3*. Put together, these results suggested that PRMT5-mediated histone symmetric dimethylation is an important and a necessary event in the regulation of *TCF3* splicing.

We further analyzed the *TCF3*-ICR region to understand if there were any peculiarities which would contribute to splicing regulation of *TCF3* involving exon 18A and 18B. We observed a CpG island to be present at the *TCF3*-ICR region (Fig. S4D) suggesting that splicing regulation could possibly be via DNA methylation. Previously, various studies have shown that DNA methylation at the splicing locus can affect alternative splicing outcomes(Pandey et al., 2024; Shukla et al., 2011; Singh et al., 2017; Yadav et al., 2023). Therefore, we first performed MeDIP-qPCR analysis in both MCF7 and MDA-MB-231 cells to understand the DNA methylation pattern in hypoxia at the *TCF3*-ICR region. We observed a significant increase in DNA methylation at the *TCF3*-ICR region under hypoxia (Fig. 5F, S4E) indicating that PRMT5-mediated symmetric arginine dimethylation of histones was somehow responsible for subsequent DNA methylation at the *TCF3*-ICR region. Next, we wanted to understand how PRMT5-mediated histone methylation was responsible for DNA methylation at the ICR region. A previously published study reported that PRMT5-mediated symmetric arginine dimethylation of histone H4 at R3 position (H4R3me2s) was capable of recruiting DNMT3A, subsequently methylating the DNA to regulate gene expression(Zhao et al., 2009). However, there were no reports suggesting the involvement of such mechanism in the regulation of alternative splicing. Therefore, we asked if histone methylation mediated DNA methylation at the *TCF3*-ICR region could possible regulate *TCF3* splicing event. Since we observed an enrichment of H4R3me2s mark at the *TCF3*-ICR along with increased DNA methylation, we wanted to check if PRMT5-mediated histone arginine symmetric dimethylation of H4R3 (H4R3me2s) could recruit DNMT3A and subsequently methylate the DNA. To check this, we first performed DNMT3A ChIP-qPCR analysis in both MCF7 and MDA-MB-231 cells under hypoxia with or without GSK591 treatment to simultaneously establish the importance of PRMT5-mediated histone arginine symmetric dimethylation in recruitment of DNMT3A to the target locus. We found a significant enrichment of DNMT3A at the *TCF3*-ICR region under hypoxia which was significantly downregulated upon GSK591 treatment (Fig. 5G, S4F). In correspondence to the decrease in DNMT3A recruitment upon GSK591 treatment, we also observed a significant downregulation in the DNA methylation at *TCF3*-ICR region upon ablation of PRMT5 methyl transferase activity (Fig. 5H, S4G) indicating that under hypoxia, PRMT5-mediated H4R3me2s mark recruits DNMT3A to the *TCF3* splicing locus and is responsible for DNA methylation. Together, these results show that PRMT5 symmetrically dimethylates H4R3 under hypoxia which in turn recruits DNMT3A which further facilitates DNA methylation of the *TCF3*-ICR region. We also observed that abrogation of the histone mark (H4R3me2s) is sufficient to inhibit DNMT3A recruitment and DNA methylation at the *TCF3*-ICR region. In summary, we established that PRMT5-mediated histone methylation at the *TCF3*-ICR region recruits DNMT3A and methylates the DNA thereby inhibiting the inclusion of exon 18A under hypoxia.

DNA methylation at the splicing locus attracts DNA binding proteins resulting in altered RNA Pol II kinetics causing differential alternative splicing outcomes (Shukla et al., 2011; Singh et al., 2017). Methyl-CpG Binding Protein 2 (MECP2) preferentially binds to methylated DNA sequences and is known to dictate alternative splicing outcomes(Brito et al., 2020; Maunakea et al., 2013). Since we observed an increase in DNA methylation at the *TCF3*-ICR region we wanted to check if MECP2 was able to bind at the *TCF3*-ICR region and regulate *TCF3* alternative splicing event under hypoxia. Hence, we performed MECP2 ChIP-qPCR analysis with or without GSK591 treatment under hypoxia to check if there was any MECP2 binding as a result of enriched DNA methylation at the *TCF3*-ICR. Certainly, we observed a significant enrichment of MECP2 at the *TCF3*-ICR region under hypoxia which was abolished upon treatment with the PRMT5 inhibitor GSK591 (Fig. 5I, S4H). Collectively, these results indicated that PRMT5-mediated H4R3me2s modification at the *TCF3*-ICR region recruited DNMT3A which subsequently led to DNA methylation at the *TCF3*-ICR region under hypoxia. Binding of MECP2 to the methylated DNA at the *TCF3*-ICR and somehow resulted in the exclusion of exon 18A under hypoxia. In order to further validate the role of MECP2 in regulating *TCF3* splicing, we performed shRNA mediated MECP2 knockdown under hypoxia (Fig. S4I) and observed changes in *TCF3* splicing pattern. Our qRT-PCR results showed a significant increase in exon 18A/18B ratio upon knocking down MECP2 under hypoxia (Fig. S5J) indicating that MECP2 somehow inhibits exon 18A inclusion under hypoxia and loss of MECP2 is sufficient to reverse the alternative splicing event.

### MECP2 binding at the methylated *TCF3*-ICR region affects RNA Pol II kinetics under hypoxia

As mentioned earlier, intragenic DNA methylation can affect RNA Pol II elongation rates affecting splicing decisions. Since we demonstrated that MECP2 binds to the methylated *TCF3*-ICR region, we suspected if it could lead to RNA Pol II pause, thereby affecting *TCF3* splicing. To check this, we performed RNA Pol II ChIP-qPCR analysis with or without GSK591 treatment in both MCF7 and MDA-MB-231 cell lines under hypoxia to understand RNA Pol II occupancy levels at the *TCF3*-ICR region and its dependence on PRMT5-mediated histone arginine symmetric dimethylation. We observed a significant increase in RNA Pol II occupancy at the *TCF3*-ICR region under hypoxia which decreased upon GSK591 treatment in both MCF7 and MDA-MB-231 cell lines (Fig. 6A). These results suggested that binding of MECP2 at the methylated *TCF3*-ICR region affected RNA Pol II kinetics and that the first step of PRMT5-mediated histone methylation is critical in initiating the cascade of events which follow. Further, to verify that increase in RNA Pol II occupancy at the *TCF3*-ICR is dependent on MECP2, we performed shRNA mediated knockdown of MECP2 in MCF7 cells under hypoxia followed by RNA Pol II ChIP-qPCR. Our ChIP analysis showed a significant reduction in RNA Pol II occupancy at the *TCF3*-ICR region upon MECP2 KD under hypoxia (Fig. 6B) which suggested that RNA Pol II elongation rate is affected by MECP2 binding and could be a critical factor in orchestrating the *TCF3* splicing event.

**Fig 6.**
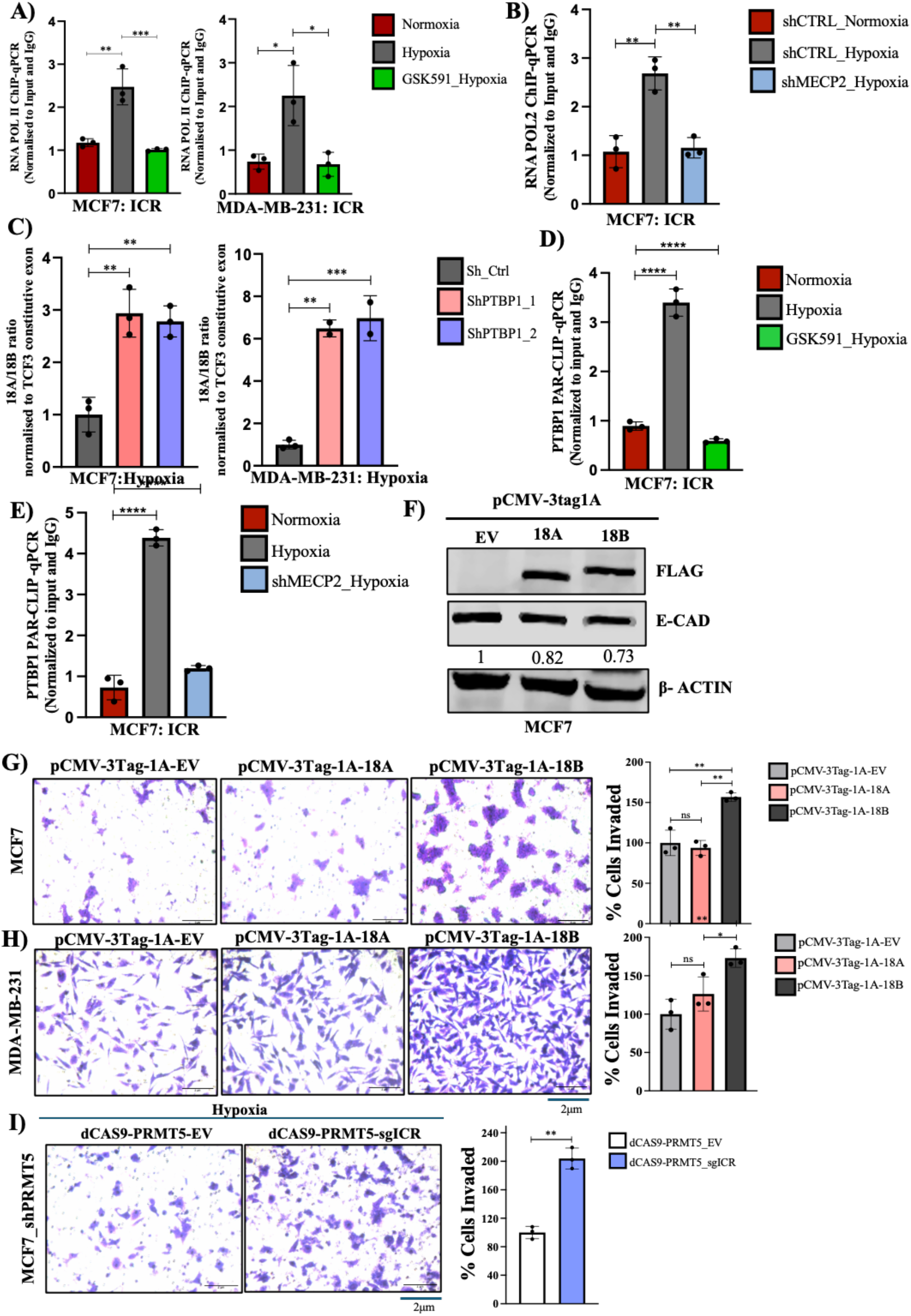
A) RNA pol II ChIP qPCR demonstrating change in RNA pol II occupancy at TCF3-ICR region in MCF7 and MDA-MB-231 cells treated with DMSO (Nx vs Hx) or 5μM GSK591(Hx). B) RNA pol II ChIP qPCR showing change in RNA pol II occupancy at TCF3-ICR region in shCTRL (Nx vs Hx) and shMECP2 (Hx) MCF7 cells. C) qRT-PCR showing increase in exon 18A/18B ratio in shPTBP1, MCF7 and MDA-MB-231 cells under hypoxia. D) PTBP1 PAR-CLIP qPCR showing change in PTBP1 binding on RNA of *TCF3* at ICR region in MCF7 cells after treatment with DMSO vs 5μM GSK591under hypoxia. E) PTBP1 PAR-CLIP qPCR showing change in PTBP1 binding on RNA of *TCF3* at ICR region in shCTRL (Nx & Hx) vs shMECP2 (Hx) MCF7 cells. F) Immunoblot showing reduction in E-Cad expression upon TCF3-18B isoform overexpression in MCF7 cells. G) Matrigel invasion assay and its respective quantification in pCMV-3tag-1A-EV vs TCF3-18A vs TCF3-18B overexpression in MCF7 cells. H) Matrigel invasion assay and its respective quantification in pCMV-3tag-1A-EV vs TCF3-18A vs TCF3-18B overexpression in MDA-MB-231 cells. I) Matrigel invasion assay and its respective quantification in shPRMT5 MCF7 cells upon transfection of dCAS9-PRMT5-EV or dCAS9-PRMT5-sgICR under hypoxia. Scale bar, 2 μm. Error bars, mean ± SEM; two-tailed t test, one way ANOVA. ∗*p* < 0.05, ∗∗*p* < 0.01, ∗∗∗*p* < 0.001, n = 3 biological replicates.

In summary, we so far demonstrated that the intronic conserved region (ICR) present between exons 18A and 18B serves as a site of control for *TCF3* splicing event. PRMT5-mediated H4R3me2s modification at the *TCF3*-ICR region increases under hypoxia which then recruits DNMT3A. Recruitment of DNMT3A subsequently results in increased DNA methylation at the *TCF3*-ICR region which serves as a binding site for MECP2. Binding of MECP2 to the methylated DNA results in slower RNA Pol II elongation rate which then becomes a critical factor in determining the *TCF3* splicing event.

### Increased RNA Pol II occupancy at the *TCF3*-ICR region recruits PTBP1 to affect *TCF3* splicing under hypoxia

Previously, it was shown that slowing of RNA Pol II could recruit a negative splicing factor, causing the exclusion of the upstream exon (Dujardin et al., 2014). Our observations so far suggested that under hypoxia, PRMT5-mediated symmetric arginine dimethylation of histones and subsequent DNA methylation, along with RNA Pol II pausing at the *TCF3*-ICR region, led to the exclusion of the upstream 18A exon. Therefore, we investigated whether RNA Pol II pausing at the *TCF3*-ICR region could possibly recruit a negative splicing factor, which could lead to the exclusion of exon 18A under hypoxia. To check this, we first performed a motif analysis of the *TCF3*-ICR region and found PTBP1 to have the highest binding score (Fig. S5A). An earlier study had revealed that PTBP1 binds to the ICR region and plays a crucial role in regulating *TCF3* splicing(Yamazaki et al., 2019), further validating our motif analysis observation. Therefore, we questioned whether PTBP1 could be the negative splicing factor responsible for exon 18A exclusion under hypoxia. To verify the involvement of PTBP1 in governing *TCF3* splicing, we performed shRNA-mediated knockdown of PTBP1 under hypoxia in both MCF7 and MDA-MB-231 cell lines (Fig. S5B) and checked if it affected *TCF3* splicing. Our qRT-PCR analysis revealed a significant increase in the exon 18A/18B ratio upon PTBP1 knockdown under hypoxia (Fig. 6C), indicating that the loss of PTBP1 resulted in increased exon 18A inclusion. This suggested that PTBP1 is a critical regulator of *TCF3* splicing which mediates exon 18A exclusion under hypoxia. To establish that PTBP1 was indeed recruited to the *TCF3*-ICR region at the RNA, we performed PAR-CLIP-qPCR analysis under hypoxia with or without GSK591 treatment in MCF7 cell line. We observed a significant enrichment of PTBP1 binding on *TCF3* RNA at the *TCF3*-ICR region, which was decreased upon GSK591 treatment (Fig. 6D, S5C), suggesting that PTBP1 recruitment was also dependent on PRMT5-mediated histone arginine symmetric dimethylation. Having hypothesized that RNA Pol II pausing allows PTBP1 recruitment to *TCF3* pre-mRNA, our next objective was to establish the significance of RNA Pol II pausing in this process. Since we had established that MECP2 binding at the *TCF3*-ICR region resulted in increased RNA Pol II occupancy, we performed shRNA mediated knockdown of MECP2 in MCF7 cell line followed by PTBP1 PAR-CLIP-qPCR analysis to observe its subsequent effect on PTBP1 recruitment. Our PAR-CLIP-qPCR analysis revealed a significant decrease in PTBP1 enrichment on *TCF3* RNA at the *TCF3*-ICR region in MCF7 cells under hypoxia (Fig. 6E) indicating that RNA Pol II pausing directly regulates PTBP1 recruitment to the *TCF3*-ICR splicing locus under hypoxia. Put together, these observations demonstrated that MECP2 mediated RNA Pol II pausing was a critical step which facilitated PTBP1 recruitment to the *TCF3* splicing locus thereafter affecting *TCF3* splicing event under hypoxia.

### Alternative splicing of TCF3 promotes EMT and invasion of breast cancer cells

Having established that PRMT5 regulates *TCF3* alternative splicing under hypoxia, we next wanted to study the importance of the alternative splicing event and its role in regulation of EMT and invasion. Under hypoxia, we observed that PRMT5 promotes exclusion of exon 18A and thereby leading to the production of the pro-invasive TCF3-18B (E47) isoform. TCF3-18B (E47) isoform is known to inhibit E-CAD expression promoting cell invasion. However, there have been no reports of TCF3 mediated regulation of invasion under hypoxia. To address this, we first cloned both and TCF3 isoforms, exon 18A or 18B into pCMV-3tag-1A FLAG-tag vector (Fig. S5D). Since normoxia mimics low TCF3-18B (E47) isoform condition, we overexpressed TCF3-18A (E12) and TCF3-18B (E47) isoforms independently in both MCF7 cell line and observed the change in E-CAD protein expression. Our immunoblot assay revealed a significant decrease in E-CAD protein levels upon TCF3-18B (E47) isoform over expression (Fig. 6F) indicating that TCF3-18B (E47) isoform induces EMT. Further to check the effect of E47 isoform on the invasion of breast cancer cells, we over expressed FLAG-tagged TCF3-18A (E12) and TCF3-18B (E47) isoforms and performed Matrigel invasion assay in both MCF7 and MDA-MB-231 cell lines. We observed that over expression of TCF3-18B (E47) isoform led to a significant increase in invasive potential of both MCF7 and MDA-MB-231 cells indicating that E47 isoform of TCF3 promotes tumor cell invasion (Fig. 6G, H, S5E). Together, these results suggested that PRMT5-mediated regulation of *TCF3* alternative splicing under hypoxia, led to the production of E47 isoform which essentially promoted EMT and invasion under hypoxia. Since we had established that PRMT5-mediated H4R3me2s mark is sufficient to regulate *TCF3* splicing which subsequently regulates tumor cell invasion, we asked if targeting dCAS9-PRMT5 to the *TCF3* splicing locus will be sufficient to enhance invasion. Therefore, we performed shRNA mediated PRMT5 KD under hypoxia followed by transfection with dCAS9-PRMT5 targeted at the TCF3-ICR splicing locus. We observed a significant increase in cell invasion upon transfection with dCAS9-PRMT5 (Fig. 6H) indicating that PRMT5-mediated histone modification at the *TCF3*-ICR splicing locus is sufficient to induce cell invasion serving as an indirect indicative of the importance of histone methylation in regulating *TCF3* alternative splicing under hypoxia.

## Discussion

Epigenetic landscape alteration and aberrant alternative splicing aid in regulation of important tumorigenic events like EMT and invasion. Epigenetic changes offer flexibility and heritability to cancer cells to weather the mounting demands of a repugnant tumor microenvironmental condition such as hypoxia. Also, alternative splicing offers transcript diversity and novelty in dealing with harsh conditions like hypoxia. Since alternative splicing event is co-transcriptional, several studies have explored the role of epigenetic features at the local transcriptional locus in dictating alternative splicing outcomes. However, the role of microenvironmental stress conditions like hypoxia in regulating epigenetics mediated changes in alternative splicing has been significantly less explored.

PRMT5 is known to aid tumor progression, yet its role and regulation in hypoxia remains underexplored. PRMT5 has been known to regulate gene expression and alternative splicing via its ability to methylate histones and spliceosome complex proteins independently. However, the ability of PRMT5 in regulating alternative splicing via modulation of histone methylation under hypoxia was not yet studied.

In this study, we systematically deduced that PRMT5 is upregulated in breast cancer patients as well as breast cancer cell lines. CCCTC binding factor (CTCF) binds at the demethylated *PRMT5* promoter under hypoxia and mediates transcriptional upregulation of *PRMT5*. PRMT5 upregulation concordantly results in an increase in PRMT5-mediated symmetric arginine dimethylation of histones H3R8 and H4R3 (H3R8me2s/H4R3me2s). We then assessed the role of PRMT5 in regulating EMT and invasion under hypoxia. We observed a PRMT5 dependent induction of EMT and tumor cell invasion under hypoxia. Subsequently, we demonstrated that PRMT5-mediated symmetric arginine dimethylation is important in regulation of EMT induction and invasion under hypoxia as inhibition of PRMT5 methyltransferase activity using GSK591 was sufficient to curb EMT and invasion under hypoxia.

Our aim was to investigate the influence of epigenetic modifications in dictating alternative splicing outcomes under hypoxia. Since PRMT5 catalyzes symmetric arginine dimethylation of histones, we hypothesized that PRMT5-mediated histone methylation regulated alternative splicing under hypoxia. To study this, we performed RNA-Seq analysis and found that under hypoxia, PRMT5 majorly regulated cassette exon events. Under hypoxia, PRMT5 promoted exon exclusion, and its absence led to an increase in exon inclusion events. Our gene ontology analysis of the genes undergoing PRMT5 dependent cassette exon splicing revealed regulation of apical junction and epithelial to mesenchymal transition (EMT) as one of the top processes regulated by PRMT5 further indicating that PRMT5 regulated EMT and invasion under hypoxia via regulation of alternative splicing.

Among all the transcription factors responsible for inducing EMT, TCF3, a bHLH domain containing transcription factor was identified as one of the novel targets of PRMT5-mediated alternative splicing. The 18A and 18B exons of *TCF3* undergo alternative splicing to give rise to either TCF3-18A (E12) isoform or TCF3-18B isoform (E47). TCF3-18B isoform inhibits E-CAD expression thereby promoting EMT and invasion. We observed a significant decrease in exon 18A/18B ration upon ablation of PRMT5 expression. Also, breast cancer cells treated with GSK591 showed decreased TCF3 exon 18A/18B ratio under hypoxia which indicated that PRMT5-mediated symmetric dimethylation is important for regulation of *TCF3* alternative splicing event. Our further analysis revealed the presence of an intronic conserved region (ICR) present between exon 18A and exon 18B of *TCF3* gene which was a potential site of regulation.

At the intronic conserved site, we observed an increase in PRMT5-mediated symmetric arginine dimethylation marks H3R8me2s and H4R3me2s under hypoxia. Using the dCAS9-PRMT5 epigenome editing system, we showed that these marks were necessary and sufficient for TCF3 alternative splicing regulation under hypoxia. PRMT5-mediated increase in H4R3me2s mark recruited DNMT3A followed by subsequent DNA methylation at the ICR under hypoxia. Methylated DNA was recognized and bound by MECP2 which resulted in RNA Pol II pausing and subsequent recruitment of the negative splicing factor PTBP1 onto the RNA at the *TCF3*-ICR. Under hypoxia, PTBP1 led to the exclusion of exon 18A leading to the production of the pro-invasive TCF3-18B (E47) isoform. Our mechanistic insights into *TCF3* splicing also reveal the intimate connection between histone marks, DNA methylation and splicing factor recruitment at the splicing locus throwing light on the innate complexity of RNA splicing. In summary, we demonstrated that PRMT5 is upregulated and promoted EMT and invasion under hypoxia in breast cancer. PRMT5-mediated symmetric arginine dimethylation of H4R3 (H4R3me2s) at the *TCF3* splicing locus is responsible for DNMT3A-MECP2-PTBP1 mediated regulation of TCF3 alternative splicing leading to the production of a pro-invasive TCF3-18B isoform which thereby promotes EMT in breast cancer (Fig. 7).

**Fig 7.**
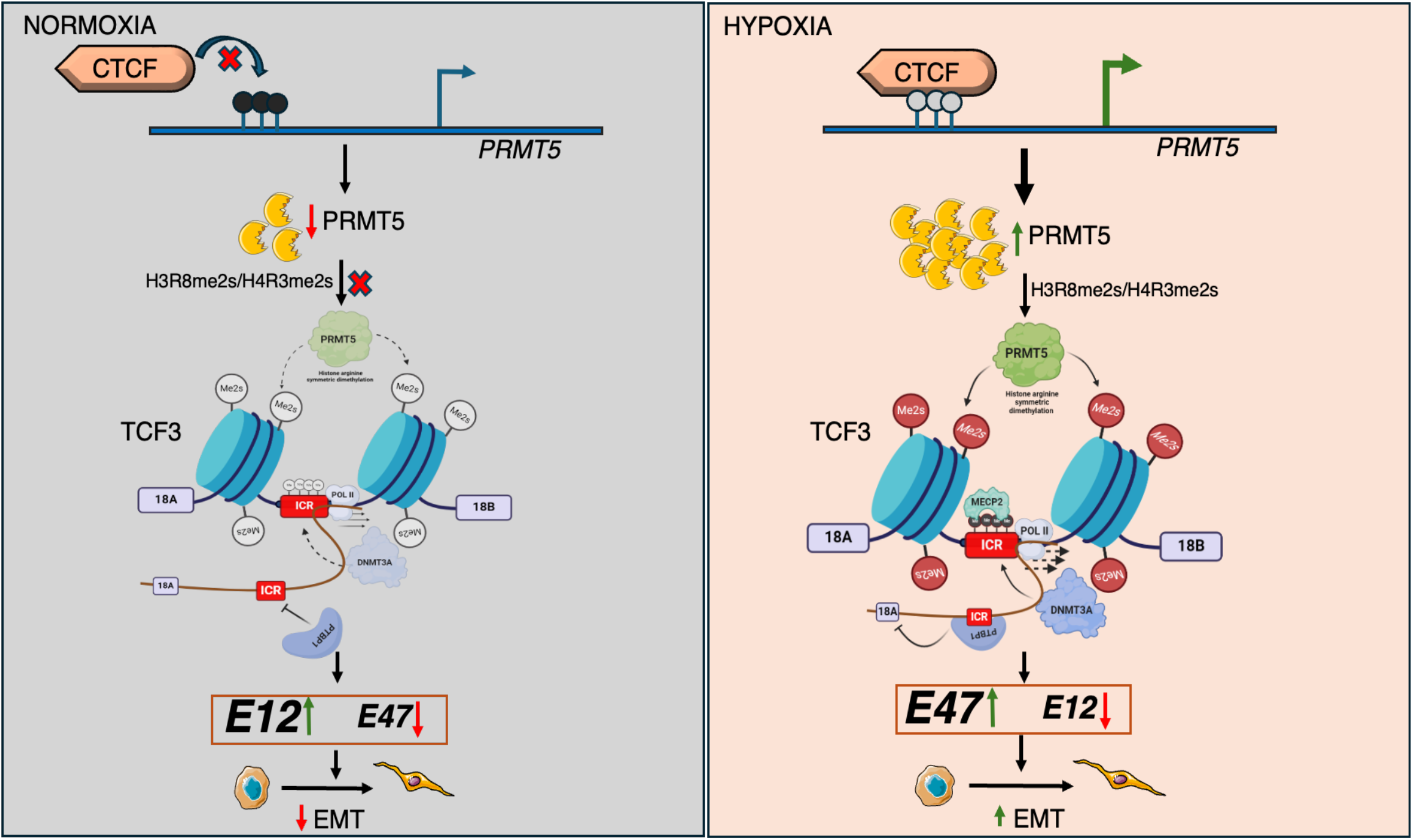
Schematic model depicting the role of PRMT5-mediated symmetric arginine dimethylation in regulation of TCF3 alternative splicing under hypoxia.

## Material and Methods

### Cell culture

Human breast cancer cell lines MCF7 and MDA-MB-231 were obtained from American Type Culture Collection (ATCC). BC8322 primary cell line derived from breast cancer patient as described in (PKM2 poised) was used in this study. MCF7, MDA-MB-231, BC8322 and HEK293T were cultured at 37ºC, 5% CO_2_ in DMEM (Invitrogen, 12800017, lot no. 2248833) supplemented with 10% fetal bovine serum (FBS; Sigma, F7524, lot no. BCBX8466), 100 units/ml of penicillin and streptomycin (Invitrogen, 15140122, lot no. 2321120), and 2 mM/l L-glutamine (Invitrogen, 25030081, lot no. 1917006). The cell lines used in the study were routinely tested for mycoplasma contamination using PCR based method. For GSK591 (Cayman Chemical, 18354) treatment, 5μM GSK591dissolved in DMSO was added to media. Cells were maintained at 1% O_2_ for hypoxia treatment in a Ruskinn INVIVO2 hypoxia chamber.

### Plasmids

The PRMT5 overexpression construct was generated by cloning the insert between *BamHI* and *EcoR*I sites of the plasmid pCMV-3tag-1a (Agilent, 240195). TCF3-18A and TCF3-18B overexpression constructs were cloned into pCMV-3tag-1a using *BamHI* and *HindIII*. For creating the dCAS9-PRMT5 construct, firstly, the DNMT3A CD was digested out of the dCAS9-DNMT3A-puroV2 plasmid (Addgene, 74407) using *BamHI* and *FseI*. Following this, MCS taken from pCMV-3tag1A was annealed and cloned in place of DNMT3A CD to create a base parent vector named as dCAS9-MCS. Subsequently, PRMT5 full length construct was cloned into the dCAS9-MCS using *BamHI* and *Sal1* to generate the dCAS9-PRMT5 construct. All of the constructs generated have been sequence verified using Sanger sequencing.

### dCAS9 mediated epigenome editing

Guide RNAs (sgRNA) were annealed and cloned at the BbsI restriction site downstream of the U6 promoter in dCAS9-DNMT3A-PuroR_v2 (Addgene, 74407)(PK paper/repurposing) or dCAS9-PRMT5 vector (This paper). Non-targeting sgRNA containing dCAS9-DNMT3A vector (Addgene, 71830) was used along with dCAS9-DNMT3A system as control. Transfections were performed in 60mm or 100mm dishes using TurboFect (Thermo scientific, R0531). For dCAS9-DNMT3A mediated epigenetic editing studies, cells were subjected to hypoxia, 12 hours post transfection. Following hypoxia treatment, cells were harvested for either immunoblot assay, MeDIP or ChIP analysis. For dCAS9-PRMT5-mediated epigenetic editing, cells were harvested 24 hours post transfection for either ChIP qPCR analysis or RNA was collected for q-RT PCR analysis.

### Quantitative RT-PCR

Total RNA was extracted using TRIzol reagent (Ambion, 15596018, lot no. 260712) as per the manufacturer’s instructions. cDNA was synthesized using PrimeScript 1st strand cDNA Synthesis Kit (Takara, 6110A). Quantitative RT-PCR (qRT-PCR) reactions were setup using KAPA SYBR^®^ FAST QPCR master mix (Sigma, KK4618) in a qTOWER^3^-G(Analytik Jena) qPCR machine according to the manufacturer’s protocol. The primers were designed using the IDT Primer Quest tool (https://www.idtdna.com) and are enlisted in Supplementary table S1. Housekeeping control gene *RPS16* was used to normalize using the formula 2^(Ct_control − Ct_target)^(Livak & Schmittgen, 2001)Student’s *t*-test or one-way ANOVA was used to compare gene/exon expression between two and three different groups respectively. *P* < 0.05 was considered statistically significant.

### Immunoblotting

Urea lysis buffer (8 M urea, 2 M thiourea, 2% CHAPS, 1% DTT) supplemented with 1× protease inhibitor cocktail (PIC; leupeptin 10–100 μM, pepstatin 1 μM, EDTA 1–10 mM, AEBSF < 1 mM) and 10mM PMSF was used to lyse the cells. Equal concentration of protein samples were loaded for every experiment. Recommended dilutions of primary antibodies was used to incubate blots overnight at 4°C, followed by 1 hour incubation with secondary antibody. The blots were scanned using Odyssey membrane Scanning system. Quantification of the bands was performed using ImageJ.

### PAR-CLIP q-RT PCR

PAR-CLIP was performed as described earlier (Spitzer et al., 2014) with slight modifications. 100 [M 4SU (4-thiouridine, Cayman Chemical, T4509) was added in the culture medium 14 hrs prior to crosslinking. Cells were crosslinked at 0.15 J^cm−2^ total energy of 365-nm UV light using a UV-crosslinker. Cells were scraped in 3mL PBS and were pelleted by centrifugation at 500Xg for 5 minutes at 4°C. Following this, cells were resuspended in NP40 lysis buffer (50mM HEPES-KOH, pH 7.5, 150mM KCl, 2mM EDTA-NaOH, pH 8, 1mM NaF, 0.5% v/v NP40) supplemented with 1X PIC, 10mM PMSF, 0.5mM DTT and 40U/mL RNaseOUT^□^ (Invitrogen, 10777019). Cells were allowed to lyse in ice for 10 minutes and were cleared at 13000 x g for 15 minutes at 4°C. Lysate was incubated with PTBP1, or rabbit IgG antibodies conjugated to Dynabeads Protein G (Invitrogen, Cat No. 10004D) overnight at 4°C. Following antibody incubation, immunoprecipitate complex was washed with IP wash buffer (50mM HEPES-KOH, pH 7.5, 300mM KCl, 0.05% v/v NP40, 0.5mM DTT (fresh), 1X PIC, 10mM PMSF, 40U/mL RNaseOUT^□^) and high salt wash buffer (50mM HEPES-KOH, pH 7.5, 500mM KCl, 0.05% v/v NP40, 0.5mM DTT (fresh)), 1X PIC, 10Mm PMSF, 40U/mL RNaseOUT^□^). 20[L aliquots of the immunoprecipitate complex were taken for immunoblot analysis. The remaining volume was subjected to decrosslinking in Proteinase K buffer (100mM Tris–HCl, pH 7.5, 12.5mM EDTA-NaOH, pH 8, 150mM NaCl, 2% v/v SDS, 1.2mg mL^-1^ Proteinase K) at 55°C for 30 minutes. TRIzol was added to the mixture and RNA was isolated as per the manufacturer’s instructions including 1 mL of Glycoblue^□^ Coprecipitant (Invitrogen, AM9515). RNA obtained was converted to cDNA with random hexamers using PrimeScript 1st strand cDNA Synthesis Kit (Takara, 6110A). qRT-PCR was performed using KAPA SYBR^®^ FAST QPCR master mix (Roche, KK4618). The enrichment of IP was normalized to its respective input using (2^(Ct IP-Ct Input). Fold change was calculated relative to IgG enrichment for representation.

### Chromatin immunoprecipitation (ChIP) assay

ChIP assay was performed as previously described (Pandkar et al., 2023). Briefly, about 10 million cells were crosslinked using 1% formaldehyde and scraped in PBS, followed by lysis and sonication. About 10-25 μg of sheared chromatin was incubated with an antibody of interest overnight at 4°C. Immunoprecipitation was performed using Dynabeads Protein G (Invitrogen, Cat No. 10004D). The immunoprecipitated protein–DNA complexes and 5% input were analysed by qRT-PCR using KAPA SYBR^®^ FAST master mix (Sigma, KK4618). Normalization was performed using the formula [2^(Ct input – Ct immunoprecipitation)^]. The obtained values were further normalized to the relative rabbit IgG and control IP values. Significance between the two groups was calculated using Student’s t-test or one-way ANOVA with a *p* value of <0.05 considered statistically significant.

### Immunohistochemistry-Fluorescence (IHC-F)

Breast tumor tissues of 5 microns size were obtained from Bansal Hospital, Bhopal, India for immunohistochemical analysis. Briefly, the sections were deparaffinized at 65°C followed by xylene treatment and were rehydrated using ethanol gradients. Heat induced antigen retrieval was performed in citrate buffer (pH 6.0) for 10 min. Blocking was done using 5% BSA for 1 hour at RT. Primary antibodies against PRMT5 and CA9 were incubated overnight at 4°C. Subsequently, tumor sections were incubated with Alexa-Flour 555 anti-rabbit IgG secondary antibody for 1 hour following which nuclear counter stain was performed using DAPI (Invitrogen, D1306, lot no. 1673432). Imaging was performed using Olympus FV3000 confocal laser scanning microscope with a 40× objective. Image analysis was performed using ImageJ software. The details of patient samples are listed in supplementary table 3

### Luciferase reporter assay

PRMT5 promoter fragments 500, 1000, 1500 and 2000 bp upstream of TSS were amplified individually. Genomic DNA obtained from MCF7 served as a template. Amplified fragments were then cloned between KpnI and HindIII restriction sites of the pGL3 basic expression vector (Promega, E1751). MCF7 or MDA-MB231 cells were seeded in 24 well plates and were co-transfected with different PRMT5 promoter luciferase constructs along with pRL-TK renilla luciferase plasmid (Promega, E2231) using PEI (Polyethyleneimine, 23966-100, Polysciences). 16 h post-transfection, the cells were subjected to appropriate hypoxia treatment. Subsequently, the cells were lysed, and the luciferase activity was determined using GloMax-Multi Detection System (Promega) by normalizing to the Renilla luciferase activity. Primers used for amplification of different fragments are listed in supplementary table 1.

### Site directed mutagenesis

To generate CTCF binding site mutant in the PRMT5 promoter luciferase construct, SDM primers containing mutations as mentioned in supplementary table S1 were used. PRMT5 -1500 promoter construct was used as a template. Following *DpnI* digestion and transformation, positive colonies were screened and the presence of intended mutations in the CTCF binding site were confirmed by Sanger sequencing.

### Matrigel invasion assay

Transwell® with 8.0 μm pore Polycarbonate membrane insert was first coated with Matrigel (Corning, 356230, lot no. 2010001). MCF7 or MDA-MB-231 cells (3 × 10^4^) were resuspended in serum free media and were added onto the Matrigel layer of the insert. The insert was placed in a well containing media supplemented with 10% FBS. The whole set up was placed in a hypoxia incubator for 24-72 hours. Following hypoxia incubation, cells were fixed with 4% formaldehyde and stained with crystal violet (0.05% in 10% methanol in 1X PBS). Images were taken at 20X in an inverted microscope (Olympus CKX41) and image analysis was performed using image j.

### RNA interference

The lentivirus containing small hairpin RNA (shRNA) (Sigma, Mission Human Genome shRNA Library) against target genes along with the packaging plasmids were transfected into HEK293T cells using PEI. Lentivirus was collected 24 and 48 hours post transfection. MCF7 and MDA-MB-23 cells were seeded at a density of 3 × 10^5^ cells per well of a six-well culture plate. Lentivirus containing media obtained from HEK293T cells was used to inoculate target cells along with 8 μg/ml polybrene (Sigma, H9268, lot no. SLBH5907V). 12 hours post viral inoculation, the media was changed and the cells were selected with 1 μg/ml puromycin (Sigma, P9620, lot no. 034M4008V) for 72 hours following which cells were used for different assays as desired. The list of shRNA sequences used in this study are provided in supplementary table S2

### Methylation Dependent Immunoprecipitation (MeDIP)

Genomic DNA (gDNA) was isolated using gDNA isolation kit (Sigma, G1N70) as per manufacturer’s protocol. MeDIP was performed as described earlier ((Kakani et al., 2024)). About 3μg of gDNA was sonicated using the Bioruptor sonicator (Diagenode) to obtain a fragment size approximately ranging from 100bp-400bp. Subsequently, sonicated DNA was denatured at 95°C for 10 mins and then incubated with 5-methyl cytosine antibody and rabbit IgG antibody overnight at 4°C. 5% of denatured DNA was taken as input. Immunoprecipitation was performed using Dynabeads Protein G (Invitrogen, 10004D). Decrosslinking of the immunoprecipitated complexes was performed in TE buffer with 1% SDS and Proteinase K (Invitrogen, 25530049) overnight at 65°C. De-crosslinked DNA and input was purified using a PCR purification kit (QIAGEN, 28106). The immunoprecipitated samples along with input were analyzed by q-RT PCR using KAPA SYBR^®^ FAST (Sigma, KK4618). The enrichment was calculated using input as normalization factor and the following formula: 2^(Ct_input − Ct_Immunoprecipitation). Additionally, the resultant values were normalized by the enrichment values of normal rabbit IgG. The final values are represented as mean ± SD of triplicates. The details of primers used for MeDIP qPCR are provided in supplementary table S1.

### RNA-Seq and bioinformatic analysis

Total RNA from hypoxia treated sh*PRMT5* and shCTRL cells was isolated using PureLink RNA Mini Kit (Invitrogen, 12183025). 1 μg of total RNA was used to make cDNA libraries. Prior to library preparation, mRNA enrichment was performed using NEBNext^®^ Poly(A) mRNA Magnetic Isolation Module (E7490S). The libraries were constructed using NEBNext^®^ Ultra^™^ II RNA Library Prep Kit for Illumina^®^ (E7770S). Paired end (2 × 150bp) sequencing was performed on Illumina NovaSeqX platform (MedGenome Labs Ltd, Bengaluru, India). Raw sequencing reads were aligned to the human reference genome (hg38 assembly) with STAR Aligner (version 2.7.1a) (Dobin et al., 2013). For alternative splicing analysis, aligned reads were analyzed using rMATS 4.0.2 (Shen et al., 2014). Differential splicing analysis was performed by calculating PSI (|⨂PSI| = |PSI (shCTRL X 100) – PSI (shPRMT5 × 100) | >3%, p <0.05) as described previously (W. juan Li et al., 2023). Genes undergoing exon inclusion upon PRMT5 knock down under hypoxia have been listed in supplementary file1. Gene ontology was performed using ShinyGo 0.80 (S. X. Ge et al., 2020). List of EMT transcription factors obtained from (Debnath et al., 2021). Spearman’s correlation analysis of *PRMT5* mRNA expression with hypoxia marker genes was performed using Gene Expression Profiling Interactive Analysis 2 (GEPIA 2) using TCGA BRCA tumor dataset (Tang et al., 2019). Transcription Factor binding sites with stringency set at 500 of breast cells was obtained from ChIP Atlas (Zou et al., 2022). UCSC genome browser hg38 tracks were used for CpG island and conservation analysis.

### ChIP-seq analysis

MCF7 cells CTCF ChIP-seq in normoxic vs hypoxic conditions was obtained from GSE216843 ((Kakani et al., 2023)). Raw fastq files were trimmed by Trimmoatic (v0.39) (Bolger et al., 2014) with default parameters. Reads were uniquely aligned to GRCh38 using STAR (v2.7.3a) aligner (Dobin et al., 2013). The biological replicates were merged by Samtools v1.9 (H. Li et al., 2009). Peak calling was performed using MACS v2.1.2 with default parameters(Zhang et al., 2008). Significant CTCF binding peaks were scanned at PRMT5 promoter. Integrative Genomics Viewer (IGV) was used to visualise CTCF binding peaks on PRMT5 promoter.

### Statistical analysis

Data are presented as mean ± SEM unless otherwise stated. At least three independent biological replicates have been performed for each experiment. All statistical analyses were conducted using GraphPad Prism 10 software. Statistical tests used and specific *p* values are indicated in the figure legends.

## Acknowledgements

S.A.M. is a recipient of funding from Indian Institute of Science Education and Research-Bhopal, India. P.Y. was supported by Council of Scientific and Industrial Research fellowship, India and is currently a post-doctoral researcher at BIDMC, Boston, MA, U.S.A. PA.K. is a recipient of post-doctoral fellowship from Indian Institute of Science Education and Research Bhopal, India, and Department of Biotechnology (DBT, India), Research Associate fellowship award. S.G.D is supported by Department of Biotechnology (DBT, India). PO.K is a graduate student at Indian Institute of Science Education and Research-Bhopal, India. This work is funded by a grant from (SERB) (CRG/2021/004949) and Department of Biotechnology (BT/PR44309/MED/30/2364/2021) to S.S.

## Author Contributions

S.S. and S.A.M and P.Y. conceived and designed the study. S.A.M., performed ChIP, immunoblotting, IHC-F, PAR-CLIP qPCR, qRT-PCR, Matrigel invasion assay, Luciferase, dCAS9 epigenetic studies, SDM, MeDIP, RNA-Seq and bioinformatic analysis. PA.K. performed CTCF ChIP and ChIP-seq analysis. P.Y. designed and cloned constructs used in Figure2. PO.K. performed q-RT PCR and immunoblotting related to Figure3. S.G.D. designed and cloned TCF3 isoform constructs. S.A.M. analysed the results. S.A.M wrote the manuscript with inputs from all other authors. S.S. supervised the experiments and manuscript preparation. S.S. acquired funding for the study.

## Data availability

RNA sequencing data from shControl and shPRMT5 cells has been deposited at GEO database (GSE279474). This study does not report any original code.

## References

Andrysik, Z., Bender, H., Galbraith, M. D., & Espinosa, J. M. (2021). Multi-omics analysis reveals contextual tumor suppressive and oncogenic gene modules within the acute hypoxic response. Nature Communications, 12(1), 1375. 10.1038/s41467-021-21687-2

Bolger, A. M., Lohse, M., & Usadel, B. (2014). Trimmomatic: a flexible trimmer for Illumina sequence data. Bioinformatics, 30(15), 2114–2120. 10.1093/bioinformatics/btu170

Brito, D. V. C., Gulmez Karaca, K., Kupke, J., Frank, L., & Oliveira, A. M. M. (2020). MeCP2 gates spatial learning-induced alternative splicing events in the mouse hippocampus. Molecular Brain, 13(1), 156. 10.1186/s13041-020-00695-1

Cao, Y., Chen, E., Wang, X., Song, J., Zhang, H., & Chen, X. (2023). An emerging master inducer and regulator for epithelial-mesenchymal transition and tumor metastasis: extracellular and intracellular ATP and its molecular functions and therapeutic potential. Cancer Cell International, 23(1), 20. 10.1186/s12935-023-02859-0

Chaffer, C. L., & Weinberg, R. A. (2011). A Perspective on Cancer Cell Metastasis. Science, 331(6024), 1559–1564. 10.1126/science.1203543

Chan-Penebre, E., Kuplast, K. G., Majer, C. R., Boriack-Sjodin, P. A., Wigle, T. J., Johnston, L. D., Rioux, N., Munchhof, M. J., Jin, L., Jacques, S. L., West, K. A., Lingaraj, T., Stickland, K., Ribich, S. A., Raimondi, A., Scott, M. P., Waters, N. J., Pollock, R. M., Smith, J. J., … Duncan, K. W. (2015). A selective inhibitor of PRMT5 with in vivo and in vitro potency in MCL models. Nature Chemical Biology, 11(6), 432–437. 10.1038/nchembio.1810

Chen, H., Lorton, B., Gupta, V., & Shechter, D. (2017). A TGFβ-PRMT5-MEP50 axis regulates cancer cell invasion through histone H3 and H4 arginine methylation coupled transcriptional activation and repression. Oncogene, 36(3), 373–386. 10.1038/onc.2016.205

Chen, Z., Han, F., Du, Y., Shi, H., & Zhou, W. (2023). Hypoxic microenvironment in cancer: molecular mechanisms and therapeutic interventions. Signal Transduction and Targeted Therapy, 8(1), 70. 10.1038/s41392-023-01332-8

Chiang, K., Zielinska, A. E., Shaaban, A. M., Sanchez-Bailon, M. P., Jarrold, J., Clarke, T. L., Zhang, J., Francis, A., Jones, L. J., Smith, S., Barbash, O., Guccione, E., Farnie, G., Smalley, M. J., & Davies, C. C. (2017). PRMT5 Is a Critical Regulator of Breast Cancer Stem Cell Function via Histone Methylation and FOXP1 Expression. Cell Reports, 21(12), 3498–3513. 10.1016/j.celrep.2017.11.096

Debnath, P., Huirem, R. S., Dutta, P., & Palchaudhuri, S. (2021). Epithelial–mesenchymal transition and its transcription factors. Bioscience Reports, 42(1), BSR20211754. 10.1042/BSR20211754

Dobin, A., Davis, C. A., Schlesinger, F., Drenkow, J., Zaleski, C., Jha, S., Batut, P., Chaisson, M., & Gingeras, T. R. (2013). STAR: ultrafast universal RNA-Seq aligner. Bioinformatics, 29(1), 15–21. 10.1093/bioinformatics/bts635

Dujardin, G., Lafaille, C., de la Mata, M., Marasco, L. E., Muñoz, M. J., Le Jossic-Corcos, C., Corcos, L., & Kornblihtt, A. R. (2014). How Slow RNA Polymerase II Elongation Favors Alternative Exon Skipping. Molecular Cell, 54(4), 683–690. 10.1016/j.molcel.2014.03.044

Duncan, K. W., Rioux, N., Boriack-Sjodin, P. A., Munchhof, M. J., Reiter, L. A., Majer, C. R., Jin, L., Johnston, L. D., Chan-Penebre, E., Kuplast, K. G., Porter Scott, M., Pollock, R. M., Waters, N. J., Smith, J. J., Moyer, M. P., Copeland, R. A., & Chesworth, R. (2016). Structure and Property Guided Design in the Identification of PRMT5 Tool Compound EPZ015666. ACS Medicinal Chemistry Letters, 7(2), 162–166. 10.1021/acsmedchemlett.5b00380

Farina, A. R., Cappabianca, L., Sebastiano, M., Zelli, V., Guadagni, S., & Mackay, A. R. (2020a). Hypoxia-induced alternative splicing: the 11th Hallmark of Cancer. Journal of Experimental & Clinical Cancer Research, 39(1), 110. 10.1186/s13046-020-01616-9

Farina, A. R., Cappabianca, L., Sebastiano, M., Zelli, V., Guadagni, S., & Mackay, A. R. (2020b). Hypoxia-induced alternative splicing: the 11th Hallmark of Cancer. Journal of Experimental & Clinical Cancer Research, 39(1), 110. 10.1186/s13046-020-01616-9

Gao, J., Liu, R., Feng, D., Huang, W., Huo, M., Zhang, J., Leng, S., Yang, Y., Yang, T., Yin, X., Teng, X., Yu, H., Yuan, B., & Wang, Y. (2021). Snail/PRMT5/NuRD complex contributes to DNA hypermethylation in cervical cancer by TET1 inhibition. Cell Death & Differentiation, 28(9), 2818–2836. 10.1038/s41418-021-00786-z

Gao, Q., Liu, Z., Liu, J., Yan, X., Dai, J., Zhang, Z., Li, R., Basnet, S., & Du, C. (2023). The role of protein arginine methylation 5 in DNA damage repair and cancer therapy. Genome Instability & Disease, 4(6), 305–314. 10.1007/s42764-023-00111-7

Ge, L., Wang, H., Xu, X., Zhou, Z., He, J., Peng, W., Du, F., Zhang, Y., Gong, A., & Xu, M. (2020). PRMT5 promotes epithelial-mesenchymal transition via EGFR-β-catenin axis in pancreatic cancer cells. Journal of Cellular and Molecular Medicine, 24(2), 1969–1979. 10.1111/jcmm.14894

Ge, S. X., Jung, D., & Yao, R. (2020). ShinyGO: a graphical gene-set enrichment tool for animals and plants. Bioinformatics, 36(8), 2628–2629. 10.1093/bioinformatics/btz931

Gupta, G. P., & Massagué, J. (2006). Cancer Metastasis: Building a Framework. Cell, 127(4), 679–695. 10.1016/j.cell.2006.11.001

Huang, J., Zheng, Y., Zheng, X., Qian, B., Yin, Q., Lu, J., & Lei, H. (2021). PRMT5 Promotes EMT Through Regulating Akt Activity in Human Lung Cancer. Cell Transplantation, 30, 09636897211001772. 10.1177/09636897211001772

Huang, Y., Hong, W., & Wei, X. (2022). The molecular mechanisms and therapeutic strategies of EMT in tumor progression and metastasis. Journal of Hematology & Oncology, 15(1), 129. 10.1186/s13045-022-01347-8

Hwang, J. W., Cho, Y., Bae, G.-U., Kim, S.-N., & Kim, Y. K. (2021). Protein arginine methyltransferases: promising targets for cancer therapy. Experimental & Molecular Medicine, 53(5), 788–808. 10.1038/s12276-021-00613-y

Imani, S., Hosseinifard, H., Cheng, J., Wei, C., & Fu, J. (2016). Prognostic Value of EMT-inducing Transcription Factors (EMT-TFs) in Metastatic Breast Cancer: A Systematic Review and Meta-analysis. Scientific Reports, 6(1), 28587. 10.1038/srep28587

Jiang, H., Zhu, Y., Zhou, Z., Xu, J., Jin, S., Xu, K., Zhang, H., Sun, Q., Wang, J., & Xu, J. (2018). PRMT5 promotes cell proliferation by inhibiting BTG2 expression via the ERK signaling pathway in hepatocellular carcinoma. Cancer Medicine, 7(3), 869–882. 10.1002/cam4.1360

Kakani, P., Dhamdhere, S. G., Pant, D., Joshi, R., Mishra, J., Samaiya, A., & Shukla, S. (2024). Hypoxia-induced CTCF promotes EMT in breast cancer. Cell Reports, 43(7). 10.1016/j.celrep.2024.114367

Kakani, P., Dhamdhere, S., Pant, D., Mishra, J., Samaiya, A., & Shukla, S. (2023). Hypoxia-induced CTCF mediates alternative splicing via coupling chromatin looping and RNA Pol II pause to promote EMT in breast cancer. BioRxiv, 2023.05.06.539689. 10.1101/2023.05.06.539689

Kim, H., & Ronai, Z. A. (2020). PRMT5 function and targeting in cancer. In Cell Stress (Vol. 4, Issue 8, pp. 199–215). Shared Science Publishers OG. 10.15698/cst2020.08.228

Li, H., Handsaker, B., Wysoker, A., Fennell, T., Ruan, J., Homer, N., Marth, G., Abecasis, G., Durbin, R., & Subgroup, 1000 Genome Project Data Processing. (2009). The Sequence Alignment/Map format and SAMtools. Bioinformatics, 25(16), 2078–2079. 10.1093/bioinformatics/btp352

Li, L., Zheng, Y.-L., Jiang, C., Fang, S., Zeng, T.-T., Zhu, Y.-H., Li, Y., Xie, D., & Guan, X.-Y. (2019). HN1L-mediated transcriptional axis AP-2γ/METTL13/TCF3-ZEB1 drives tumor growth and metastasis in hepatocellular carcinoma. Cell Death & Differentiation, 26(11), 2268–2283. 10.1038/s41418-019-0301-1

Li, W. juan, Huang, Y., Lin, Y. an, Zhang, B. ding, Li, M. Y., Zou, Y. qin, Hu, G. sheng, He, Y. hui, Yang, J. jing, Xie, B. lan, Huang, H. hua, Deng, X., & Liu, W. (2023). Targeting PRMT1-mediated SRSF1 methylation to suppress oncogenic exon inclusion events and breast tumorigenesis. Cell Reports, 42(11). 10.1016/j.celrep.2023.113385

Li, Z. X., Sun, M. C., Fang, K., Zhao, Z. Y., Leng, Z. Y., Zhang, Z. H., Xu, A. P., Chu, Y., Zhang, L., Lian, J., Chen, T., & Xu, M. D. (2023). Transcription factor 3 promotes migration and invasion potential and maintains cancer stemness by activating ID1 expression in esophageal squamous cell carcinoma. Cancer Biology and Therapy, 24(1). 10.1080/15384047.2023.2246206

Liu, F., Xu, Y., Lu, X., Hamard, P.-J., Karl, D. L., Man, N., Mookhtiar, A. K., Martinez, C., Lossos, I. S., Sun, J., & Nimer, S. D. (2020). PRMT5-mediated histone arginine methylation antagonizes transcriptional repression by polycomb complex PRC2. Nucleic Acids Research, 48(6), 2956–2968. 10.1093/nar/gkaa065

Livak, K. J., & Schmittgen, T. D. (2001). Analysis of Relative Gene Expression Data Using Real-Time Quantitative PCR and the 2-ΔΔCT Method. Methods, 25(4), 402–408. 10.1006/meth.2001.1262

López-Menéndez, C., Vázquez-Naharro, A., Santos, V., Dubus, P., Santamaría, P. G., Martínez-Ramírez, Á., Portillo, F., Moreno-Bueno, G., Faraldo, M. M., & Cano, A. (2021). E2A Modulates Stemness, Metastasis, and Therapeutic Resistance of Breast Cancer. Cancer Research, 81(17), 4529–4544. 10.1158/0008-5472.CAN-20-2685

Luco, R. F., Pan, Q., Tominaga, K., Blencowe, B. J., Pereira-Smith, O. M., & Misteli, T. (2010). Regulation of alternative splicing by histone modifications. Science, 327(5968), 996–1000. 10.1126/science.1184208

Maunakea, A. K., Chepelev, I., Cui, K., & Zhao, K. (2013). Intragenic DNA methylation modulates alternative splicing by recruiting MeCP2 to promote exon recognition. Cell Research, 23(11), 1256–1269. 10.1038/cr.2013.110

Pandey, A., Kakani, P., & Shukla, S. (2024). CTCF and BORIS-mediated autophagy regulation via alternative splicing of BNIP3L in breast cancer. Journal of Biological Chemistry, 300(7), 107416. 10.1016/j.jbc.2024.107416

Pandkar, M. R., Raveendran, A., Biswas, K., Mutnuru, S. A., Mishra, J., Samaiya, A., Malys, T., Mitr Ophano V, A. Y., Sharan, S. K., & Shukla, S. (2023). PKM2 dictates the poised chromatin state of PFKFB3 promoter to enhance breast cancer progression. 5(3). 10.1093/narcan/z

Rengasamy, M., Zhang, F., Vashisht, A., Song, W.-M., Aguilo, F., Sun, Y., Li, S., Zhang, W., Zhang, B., Wohlschlegel, J. A., & Walsh, M. J. (2017). The PRMT5/WDR77 complex regulates alternative splicing through ZNF326 in breast cancer. Nucleic Acids Research, 45(19), 11106–11120. 10.1093/nar/gkx727

Riggio, A. I., Varley, K. E., & Welm, A. L. (2021). The lingering mysteries of metastatic recurrence in breast cancer. British Journal of Cancer, 124(1), 13–26. 10.1038/s41416-020-01161-4

Sachamitr, P., Ho, J. C., Ciamponi, F. E., Ba-Alawi, W., Coutinho, F. J., Guilhamon, P., Kushida, M. M., Cavalli, F. M. G., Lee, L., Rastegar, N., Vu, V., Sánchez-Osuna, M., Coulombe-Huntington, J., Kanshin, E., Whetstone, H., Durand, M., Thibault, P., Hart, K., Mangos, M., … Dirks, P. B. (2021). PRMT5 inhibition disrupts splicing and stemness in glioblastoma. Nature Communications, 12(1), 979. 10.1038/s41467-021-21204-5

Saitoh, M. (2023). Transcriptional regulation of EMT transcription factors in cancer. Seminars in Cancer Biology, 97, 21–29. 10.1016/j.semcancer.2023.10.001

Sapir, T., Shifteh, D., Pahmer, M., Goel, S., & Maitra, R. (2021). Protein Arginine Methyltransferase 5 (PRMT5) and the ERK1/2 & PI3K Pathways: A Case for PRMT5 Inhibition and Combination Therapies in Cancer. Molecular Cancer Research, 19(3), 388–394. 10.1158/1541-7786.MCR-20-0745

Segelle, A., Núñez-Álvarez, Y., Oldfield, A. J., Webb, K. M., Voigt, P., & Luco, R. F. (2022a). Histone marks regulate the epithelial-to-mesenchymal transition via alternative splicing. Cell Reports, 38(7). 10.1016/j.celrep.2022.110357

Segelle, A., Núñez-Álvarez, Y., Oldfield, A. J., Webb, K. M., Voigt, P., & Luco, R. F. (2022b). Histone marks regulate the epithelial-to-mesenchymal transition via alternative splicing. Cell Reports, 38(7), 110357. 10.1016/j.celrep.2022.110357

Sengupta, S., West, K. O., Sanghvi, S., Laliotis, G., Agosto, L. M., Lynch, K. W., Tsichlis, P. N., Singh, H., Patrick, K. L., & Guerau-de-Arellano, M. (2021). PRMT5 Promotes Symmetric Dimethylation of RNA Processing Proteins and Modulates Activated T Cell Alternative Splicing and Ca2+/NFAT Signaling. ImmunoHorizons, 5(10), 884–897. 10.4049/immunohorizons.2100076

Shen, S., Park, J. W., Lu, Z., Lin, L., Henry, M. D., Wu, Y. N., Zhou, Q., & Xing, Y. (2014). rMATS: Robust and flexible detection of differential alternative splicing from replicate RNA-Seq data. Proceedings of the National Academy of Sciences, 111(51), E5593–E5601. 10.1073/pnas.1419161111

Shukla, S., Kavak, E., Gregory, M., Imashimizu, M., Shutinoski, B., Kashlev, M., Oberdoerffer, P., Sandberg, R., & Oberdoerffer, S. (2011). CTCF-promoted RNA polymerase II pausing links DNA methylation to splicing. Nature, 479(7371), 74–79. 10.1038/nature10442

Singh, S., Narayanan, S. P., Biswas, K., Gupta, A., Ahuja, N., Yadav, S., Panday, R. K., Samaiya, A., Sharan, S. K., & Shukla, S. (2017). Intragenic DNA methylation and BORIS-mediated cancer-specific splicing contribute to the Warburg effect. Proceedings of the National Academy of Sciences, 114(43), 11440–11445. 10.1073/pnas.1708447114

Spitzer, J., Hafner, M., Landthaler, M., Ascano, M., Farazi, T., Wardle, G., Nusbaum, J., Khorshid, M., Burger, L., Zavolan, M., & Tuschl, T. (2014). PAR-CLIP (Photoactivatable Ribonucleoside-Enhanced Crosslinking and Immunoprecipitation): A Step-By-Step Protocol to the Transcriptome-Wide Identification of Binding Sites of RNA-Binding Proteins. In Methods in Enzymology (Vol. 539, pp. 113–161). Academic Press Inc. 10.1016/B978-0-12-420120-0.00008-6

Tang, Z., Kang, B., Li, C., Chen, T., & Zhang, Z. (2019). GEPIA2: an enhanced web server for large-scale expression profiling and interactive analysis. Nucleic Acids Research, 47(W1), W556–W560. 10.1093/nar/gkz430

Yadav, P., Pandey, A., Kakani, P., Mutnuru, S. A., Samaiya, A., Mishra, J., & Shukla, S. (2023). Hypoxia-induced loss of SRSF2-dependent DNA methylation promotes CTCF-mediated alternative splicing of VEGFA in breast cancer. IScience, 26(6), 106804. 10.1016/j.isci.2023.106804

Yamazaki, T., Liu, L., Lazarev, D., Al-Zain, A., Fomin, V., Yeung, P. L., Chambers, S. M., Lu, C. W., Studer, L., & Manley, J. L. (2018). TCF3 alternative splicing controlled by hnRNP H/F regulates E-cadherin expression and hESC pluripotency. Genes and Development, 32(17–18), 1161–1174. 10.1101/gad.316984.118

Yamazaki, T., Liu, L., & Manley, J. L. (2019). TCF3 mutually exclusive alternative splicing is controlled by long-range cooperative actions between hnRNPH1 and PTBP1. 10.1261/rna

Zhang, Y., Liu, T., Meyer, C. A., Eeckhoute, J., Johnson, D. S., Bernstein, B. E., Nusbaum, C., Myers, R. M., Brown, M., Li, W., & Liu, X. S. (2008). Model-based Analysis of ChIP-Seq (MACS). Genome Biology, 9(9), R137. 10.1186/gb-2008-9-9-r137

Zhao, Q., Rank, G., Tan, Y. T., Li, H., Moritz, R. L., Simpson, R. J., Cerruti, L., Curtis, D. J., Patel, D. J., Allis, C. D., Cunningham, J. M., & Jane, S. M. (2009). PRMT5-mediated methylation of histone H4R3 recruits DNMT3A, coupling histone and DNA methylation in gene silencing. Nature Structural & Molecular Biology, 16(3), 304–311. 10.1038/nsmb.1568

Zou, Z., Ohta, T., Miura, F., & Oki, S. (2022). ChIP-Atlas 2021 update: a data-mining suite for exploring epigenomic landscapes by fully integrating ChIP-seq, ATAC-seq and Bisulfite-seq data. Nucleic Acids Research, 50(W1), W175–W182. 10.1093/nar/gkac199.

